# Nonsense-mediated mRNA decay of metal-binding activator *MAC1* is dependent on copper levels and 3′-UTR length in *Saccharomyces cerevisiae*

**DOI:** 10.1101/2022.10.03.510652

**Authors:** Xinyi Zhang, Bessie W. Kebaara

**Affiliations:** Department of Biology, One Bear Place #97388, Baylor University, Waco, TX 76798, USA

## Abstract

The nonsense-mediated mRNA decay (NMD) pathway was initially identified as a pathway that degrades mRNAs with premature termination codons but is now also recognized as a post-transcriptional gene regulatory pathway which regulates expression of natural mRNAs. Earlier studies demonstrated that regulation of functionally related mRNAs by NMD can be differential and conditional in *Saccharomyces cerevisiae*. Here, we elucidate the regulation of *MAC1* mRNAs by NMD in response to copper, and the role *MAC1* 3′-UTR plays in this regulation. *MAC1* is a copper-sensing transcription factor that regulates high affinity copper transport and is activated under low copper conditions in *S. cerevisiae*. We found that *MAC1* mRNAs were regulated by NMD under normal growth conditions but escape NMD under low and high copper conditions. Mac1p regulated genes escape NMD in conditions where *MAC1* mRNAs are NMD sensitive. We also found that the *MAC1* 3′-UTR contributes to the degradation of the mRNAs by NMD and that *MAC1* mRNAs lacking 3′-UTR were stabilized in response to copper deprivation. Taken together, our results demonstrate a novel mechanism of regulating a metal sensing transcription factor, where *MAC1* mRNA levels are regulated by NMD and copper, while the activity of Ma1p is controlled by copper levels.

## INTRODUCTION

The nonsense-mediated mRNA decay (NMD) pathway is a highly conserved mRNA degradation pathway. It was primarily discovered to be an mRNA surveillance system that protects eukaryotic cells by reducing the production of harmful truncated proteins from premature termination codon (PTC) bearing transcripts (1). The substrates of this pathway are transcripts derived from genes harboring nonsense mutations. NMD is now also recognized as a pathway that regulates natural mRNAs that encode fully functional proteins. The targeting features of some natural mRNAs include inefficiently spliced pre-mRNAs that enter the cytoplasm, mRNAs with atypical long 3′-UTR, mRNAs in which the ribosome has bypassed the initiator AUG codon and commenced translation further downstream, some mRNAs that contain upstream open reading frames (uORFs), and mRNAs that are subject to frameshifting or that are generated by certain alternative splicing events (2-7).

Regulation of natural mRNAs by NMD has been observed in diverse organisms including *Saccharomyces cerevisiae, Drosophila melanogaster, Caenorhabditis elegans*, and humans (3,8-12). In the yeast *Saccharomyces cerevisiae*, when NMD is inactivated, 5-10% of the transcriptome is affected. Hundreds of endogenous RNA Polymerase II transcripts achieve steady-state levels that depend on NMD. The decay rate of direct NMD targets is directly influenced by NMD, while abundance of indirect targets is NMD-sensitive but without any effect on the decay rate (10). From yeast to humans, activation of NMD requires the function of three principal conserved up-frameshift (UPF) factors: Upf1, Upf2, and Upf3 (13,14).

The regulation of specific natural mRNAs by the NMD results in a decreased pool of the mRNA which should result in less protein. Thus, NMD modulates the expression of select genes by precisely regulating the expression of specific mRNA. Importantly, natural mRNAs regulated by NMD contain features that trigger their degradation and occur in specific cellular context and environmental conditions (15). That is, NMD controls mRNA levels in response to environmental stimuli. Many different stimuli can induce NMD-mediated degradation including nutritional metals such as copper and iron. We found that in *S. cerevisiae* NMD plays a role in copper and iron homeostasis (16-18). Understanding nutritional metal homeostatic mechanisms in yeast will provide insights into the regulatory mechanisms used by other organisms as these mechanisms have been conserved throughout evolution (19).

The transcription factor Mac1 (<u>m</u>etal-binding <u>ac</u>tivator) in *S. cerevisiae* plays a critical role in regulating the high-affinity copper uptake system and is activated under low copper conditions (20,21). In response to low copper conditions, Mac1p binds to Cu-responsive *cis*-acting (CuRE) promoter elements of target genes and activates the high-affinity copper uptake systems encoded by *CTR1* and *CTR3* (22). Mac1p also regulates the expression of two cell-surface metal reductases encoded by *FRE1* and *FRE7* (22,23). Therefore, we would anticipate the *MAC1* mRNA would be stabilized in wild-type yeast strains under this specific growth condition when higher levels of copper are required. Our previous analysis of mRNAs encoding proteins involved in various aspects of copper homeostasis, demonstrated that regulation of these mRNAs by NMD could be responsive to fluctuations in environmental copper levels and generates physiological consequences to the yeast cells (17,24).

Here we investigated the regulation of *MAC1* mRNA by NMD under changing levels of copper. *MAC1* encodes two mRNA isoforms generated by alternative 3′-end processing. The major *MAC1* mRNA isoform is regulated by NMD in a condition specific manner. We found that this mRNA is a direct NMD target in normal growth conditions (CM) but is not regulated by the pathway in low or high copper conditions. Interestingly Mac1p regulated genes, *CTR1* and *FRE1* are insensitive to NMD under normal growth conditions or in low or high copper conditions. The alternative 3′-end processing of *MAC1* 3′-UTR is transferrable to *CYC1*, but not influenced by environmental copper conditions. Additionally, we found that the shorter *MAC1* 3′-UTR is not sufficient to target NMD insensitive *CYC1* mRNA to NMD pathway, while the longer *MAC1* 3′-UTR is sufficient to directly target the *CYC1MAC1* 3′-UTR isoform to NMD. Furthermore, the absence of the 3′-UTR from *MAC1* mRNA renders the mRNA insensitive to the NMD pathway but preserves *MAC1* mRNA sensitivity to low copper conditions. This study demonstrates that regulation of *MAC1* in response to environmental copper conditions is more complex than merely at the protein level. While *MAC1* mRNA is regulated by NMD in a condition specific manner, activity of Mac1p is copper dependent. Overall, the control of *MAC1* transcript levels by NMD and by copper is unique. Copper deficiency influences alternative polyadenylation of *MAC1* mRNA and the two *MAC1* transcripts produced are immune to NMD. Mac1 protein activity is controlled by copper levels.

## MATERIAL AND METHODS

### Yeast strains

All *S. cerevisiae* strains used in this study are listed in Supplementary Table S1. Yeast strains were grown and maintained using standard techniques. Strains used to measure mRNA half-lives have the temperature-sensitive allele of RNA polymerase II, *rpb1-1*(25). *rpb1-1* yeast strains grow at 28°C (permissive temperature) and transcription is inhibited at 39°C (the nonpermissive temperature). Subsequently, mRNA decay rate is measured at different time points after transcription has been inhibited at the nonpermissive temperature. For mRNA half-lives of yeast strains lacking the *rpb1-1* allele, the cells are treated with thiolutin to inhibit transcription. mRNA decay rate was measured at the same time points as in *rpb1-1* yeast strains after transcription was inhibited using 2mg/ml thiolutin. Typical mRNA half-lives were measured over a 35-min time period.

### Growth under high and low copper

For growth under low copper conditions, yeast strains were grown in low copper complete minimal media. To attain low copper conditions, the media contained yeast nitrogen base without copper and iron (YNB-CuSO_4_-FeCl_3_) and 100 μM Bathocuproinedisulfonic acid (BCS) (Sigma-Aldrich). Glassware used in these experiments was soaked in 10% nitric acid overnight to remove trace amounts of copper. All yeast cells used for low copper were initially grown to saturation in complete minimal media then sub-cultured into copper deficient media in acid washed glassware.

To analyze the mRNAs under high copper conditions, wild-type and NMD mutant yeast cells were grown in CM media supplemented with 600 μM copper (high copper media). As with low copper conditions, the yeast cells were first grown to saturation in complete minimal media then sub-cultured into media supplemented with 600 μM copper.

### RNA methods

For all mRNA steady-states and half-life experiments, total *S. cerevisiae* RNA was used. Yeast cells cultured in the conditions described above were harvested at mid-log phase as described in Peccarelli and Kebaara(26). Total RNA was extracted from harvested cells using the hot phenol method and run on an agarose-formaldehyde gel. The RNA was transferred to GeneScreen Plus^®^ (PerkinElmer, Boston, MA) nylon membranes using the NorthernMax^™^ Complete Northern Blotting kit (Thermo Fisher Scientific, Carlsbad, CA) transfer protocol. Northern blots were probed with oligolabeled DNA probes that were labeled with [α-^32^P] dCTP using the RadPrime DNA Labeling System (Thermo Fisher Scientific, Carlsbad, CA). All probes were generated by PCR. Northern blots were phosphorImaged^™^ using a Typhoon Phosphorimager (Amersham Pharmacia Biotech, Inc.). All northern blots were probed with *CYH2* as an NMD control to confirm the NMD phenotype of the yeast strains. *CYH2* pre-mRNA is a known NMD target, while *CYH2* mRNA is not. *CTR1* mRNA was used as low copper control. *CTR1* encodes a high-affinity copper transporter of plasma membrane, and diminished copper levels result in increased *CTR1* expression. *CUP1* mRNA was used as high copper control. *CUP1* encodes a metallothionein that binds copper, and *CUP1* mRNA is not sensitive to NMD(27). *SCR1* was used as a loading control for all northern blots. *SCR1* is an RNA polymerase III transcript that is not regulated by NMD or sensitive to copper or iron levels in the environment. All northerns were quantified using ImageQuant software. Sigmaplot 2000, Version 14.0 software was used to calculate half-lives as described in Peccarelli and Kebaara (26).

### 3′ Rapid Amplification of cDNA ends (3′-RACE)

3′-RACE was used to determine the length of the mRNA 3′UTRs using the 3′RACE System for Rapid Amplification of cDNA Ends kit (Thermo Fisher Scientific, Carlsbad, CA, USA). Yeast total RNA used for steady-state and half-life northern blots was used to generate cDNA using SuperScript^™^II RT (Thermo Fisher Scientific, Carlsbad, CA, USA). Subsequently, the cDNA was used as the template for all primary PCR reactions. Primary PCR reactions used the Abridged Universal Amplification Primer (AUAP) from the 3′RACE kit in combination with gene-specific primers. The primary PCR product served as a template for the nested PCR reactions. All nested PCR reactions utilized gene specific primers. PCR products for both primary and nested reactions were run on 1.5% agarose gels.

### DNA methods

To generate the *CYC1MAC1 3′-UTR* fusion construct, the long 3′-UTR from *MAC1* was amplified by PCR. Subsequently, DNA containing the 5′-UTR and ORF of a second gene, *CYC1*, was amplified by PCR. *CYC1* mRNA is not an NMD target. We and others have used it previously to characterize NMD-targeting features (16,17,27,28). Third, ligation-mediated PCR fused the two PCR fragments. Alternatively, to create *MAC1CYC1* 3′-UTR, DNA comprising the 5′-UTR and ORF of *MAC1* was fused to 350 nt from the *CYC1* 3′-UTR. The *MAC1CYC1* 3′-UTR construct contained the *MAC1* promoter and *CYC1* terminator sequences.

The fusion constructs were then inserted into the TOPO-TA cloning vector according to manufacturer’s instructions and sent for sequencing to verify sequences and confirm that the correct fusion construct was generated. Next, *CYC1MAC1* 3′-UTR in TOPO-TA were digested with SpeI and NotI prior to ligation into the yeast centromeric vector pRS315. The *MAC1CYC1* 3′-UTR in TOPO-TA were digested with BamHI and NotI prior to ligation into the yeast vector pRS315 (29). Lithium acetate mediated transformation was used to transform the plasmids into wild-type and NMD mutant strains. All plasmids were maintained in complete minimal media lacking leucine.

### Statistical analysis

Tests for significance were done using a two tailed t test. All experiments were done in triplicate unless otherwise stated. Significance was defined by p values determined from the t test. P < 0.05 was considered a significant difference in mRNA steady-state levels and half-life comparisons.

## RESULTS

### Regulation of *MAC1* mRNA steady-state levels by NMD is dependent on copper levels

Because Mac1p is a copper-sensing transcription factor that activates the expression of genes under low copper conditions, we examined if *MAC1* mRNAs are regulated by NMD under different environmental copper conditions. *MAC1* mRNA steady state levels were measured in wild-type and NMD mutants (*upf1Δ*) cells grown in complete minimal (CM), CM media containing 100 μM Bathocuproinedisulfonic acid (BCS) (low copper) and CM media containing 600 μM copper (high copper).

In complete minimal media, *MAC1* mRNA accumulated 1.6 (± 0.2) fold (p-value = 0.02) higher in the NMD mutant relative to the wild-type strain (Figure 1B and Table 1). Additionally, we found that *MAC1* mRNA accumulated to higher levels in low copper conditions compared to normal growth conditions (CM). In wild-type cells, *MAC1* mRNA accumulated 4.7 (±1.1) fold higher in low copper conditions relative to CM, while in NMD mutant cells, *MAC1* mRNA accumulated 3.3 (±1.1) fold higher in low copper conditions relative to CM. However, there was no significant difference in *MAC1* mRNA steady-state accumulation levels between wild-type and NMD mutant in low copper conditions. The most abundant *MAC1* mRNA isoform accumulated 1.1 (± 0.6) under low copper conditions (Figure 1B and Table.1). Similarly, the predominant *MAC1* mRNA isoform accumulated 0.9 (± 0.2) fold in NMD mutants (*upf1Δ*) under high copper conditions (600 μM copper) relative to wild-type cells (Figure 1B). These observations suggest that *MAC1* mRNA is only regulated by NMD under CM, but not in copper deplete (100 μM BCS) and copper replete conditions (600 μM copper). Furthermore, the expression levels of Mac1p activated gene *CTR1*, which encodes a plasma membrane copper transporter were elevated in low copper conditions and undetectable in copper replete conditions (Figure 1B). Interestingly, *CTR1* mRNA did not accumulate to higher levels in NMD mutants relative to wild-type strains under normal growth conditions where *MAC1* mRNA is regulated by NMD (Figure 1B).

**Figure 1.**
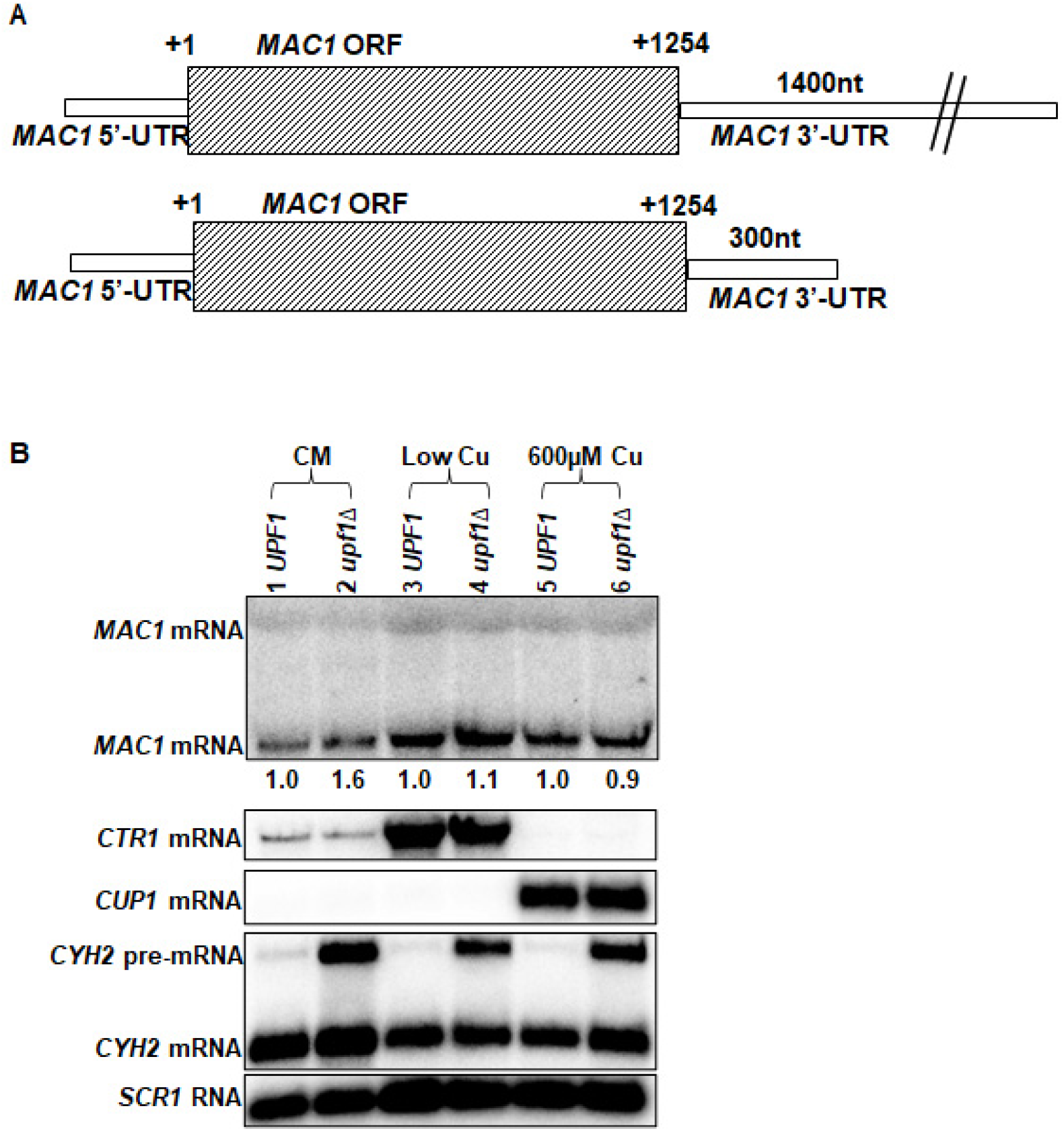
Regulation of *MAC1* mRNA steady-state levels by NMD is dependent on copper levels. Schematic representation of *MAC1* mRNA isoforms (A). Representative mRNA steady-state accumulation levels of *MAC1* mRNA (B). mRNA levels were measured with total RNA from wild-type strain W303 (*UPF1*) (39), and NMD mutants AAY320 (*upf1Δ*)(25). The northern blot was probed with DNA specific to *MAC1* open reading frame (ORF). The major *MAC1* mRNA isoform fold change (*upf1Δ/UPF1*) are shown below of the northern blot (B). *CTR1, CUP1, CYH2* and *SCR1* were used as controls. *CTR1* was used as low copper control. *CTR1* encodes a high affinity copper transporter of the plasma membrane. *CTR1* gene is activated under low copper conditions. *CUP1* was used as a control for high copper because *CUP1* encodes a metallothionein that binds copper. The *CUP1* gene is induced by the Ace1 transcription factor when cells are exposed to elevated copper levels. *CYH2* pre-mRNA was used as an NMD control because *CYH2* pre-mRNA is degraded by NMD. *SCR1* was used as a loading control for all northern blots. *SCR1* is an RNA polymerase III transcript that is not regulated by NMD.

**Table 1.** *MAC1* mRNA relative accumulation levels under different growth conditions. Wild-type (W303) and NMD mutants (AAY320) were used in mRNA steady-state accumulation measurements. mRNA steady-state accumulation levels were done in triplicate and reported as an average ± standard deviation (SD). BCS* - (Bathocuproinedisulfonic acid) Low copper.

| Growth media | Relative mRNA accumulation (*upf1Δ/UPF1*) |
| --- | --- |
| Complete minimal | 1.6 (± 0.2) |
| 100 μM BCS* | 1.1 (± 0.6) |
| 600 μM Cu | 0.9 (± 0.2) |

### *MAC1* mRNA is directly regulated by NMD in complete minimal but not under low or high copper conditions

To confirm that *MAC1* is a direct NMD target only under CM, we measured the half-life of *MAC1* mRNA in CM and low copper conditions using wild-type (*UPF1 rpb1-1)* and NMD mutant (*upf1Δ rpb1-1)* yeast strains. These strains harbor the temperature-sensitive allele of RNA polymerase II, *rpb1-1*(26). Measurement of *MAC1* mRNA half-lives in these strains showed that there was a significant difference (p-value = 0.02) in the mRNA decay rates in complete minimal media (Figure 2A and Supplementary Figure 1A). Specifically, the half-life of the major *MAC1* mRNA isoform in the wild-type strains was 5.3 (± 0.8) mins relative to 7.8 (± 0.8) mins in the NMD mutants (Figure 2A and Table 2).

**Figure 2.**
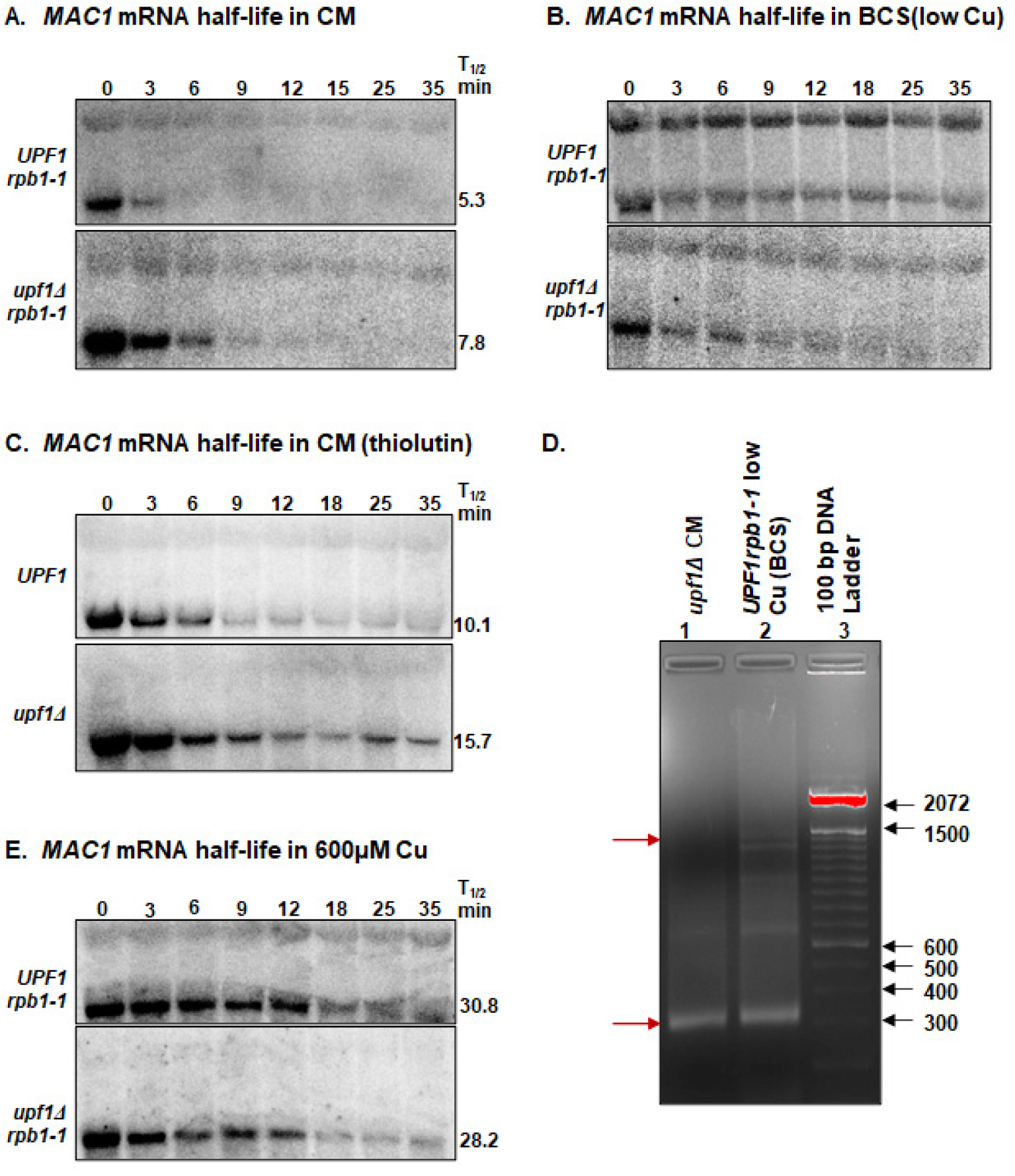
*MAC1* mRNA is directly regulated by NMD under regular growth conditions (CM) but resistant to NMD under low copper (BCS) and high copper. Representative half-life northern blots of *MAC1* mRNA. The mRNA half-lives were measured with total RNA extracted from wild-type strain AAY334 (*UPF1 rpb1-1*)(25) and NMD mutant strain AAY335 (*upf1Δ rpb1-1*) (25) grown under complete minimal (A), low copper (B) and in 600 μM Cu, high copper (E). mRNA half-lives were measured with total RNA extracted from wild-type strain W303 (*UPF1*) (39), and NMD mutants (*upf1Δ*)(25) grown under complete minimal treated with thiolutin (C). Yeast cells were harvested over a 35-min period at eight time points indicated above the northern blots. The half-lives were determined using SigmaPlot and are shown to the right of the northern blots. All half-life measurements are an average of three independent experiments. For controls, the membranes were probed with DNA specific to *CTR1, FRE1, CUP1, CYH2* and *SCR1* ORFs (Supplementary Figure 1). A 1.5% agarose gel of primary 3′-RACE PCR products of the *MAC1* mRNA isoforms in wild-type and *upf1Δ* stains grown in complete minimal or low copper medium (D). The arrows indicate the size of the 3′RACE PCR products expected to be present in the two *MAC1* mRNA isoforms. The additional bands seen on the 3′RACE PCR gel were considered non-specific bands since there were no corresponding bands on the respective northern blot.

**Table 2.** *MAC1* mRNA half-lives under different growth conditions. Wild-type (AAY334) and NMD mutants (AAY335) have the *rpb1-1* allele. All yeast strains used were grown under regular growth conditions in complete minimal media, 100 μM BCS (low copper), and 600 μM copper. W303 and AAY320 were treated with thiolutin to determine half-lives under regular conditions in complete minimal media. The half-life experiments were done in triplicate and are reported as averages ± standard deviation. ND, not determined.

| Growth media | mRNA half-life<br>(min) *UPF1* | mRNA half-life<br>(min) *upf1Δ* |
| --- | --- | --- |
| Complete minimal in *rpb1-1* | 5.3 (± 0.8) | 7.8 (± 0.8) |
| Complete minimal with thiolutin | 10.1 (± 0.4) | 15.7 (± 1.2) |
| 100 μM BCS* in *rpb1-1* | ND | ND |
| 600 μM Cu in *rpb1-1* | 30.8 (± 4.7) | 28.2 (± 6.1) |

ND- not determined

Additionally, the half-lives of the *MAC1* transcripts under low copper were measured using the same time points as the ones used to measure the half-lives in CM for comparison purposes (Figure 2B and Supplementary Figure S1B). Two major *MAC1* mRNA isoforms were detected under low copper as previously reported (15). The longer *MAC1* mRNA isoform accumulated to higher levels relative to normal growth conditions (Figure 2A and 2B). The decay rate of the major *MAC1* mRNA isoforms varied from the decay rate in complete minimal. Specifically, the *MAC1* mRNA half-life in both wild type and NMD mutant strains could not be determined using these time points because the mRNA did not decay. This is consistent with the previous study from our lab showing that *MAC1* mRNA was not degraded within the 35-minute time period (15). The relative abundance of the long and short *MAC1* mRNA isoforms at 0 min time point were significantly different in low copper conditions relative to CM. Specifically, the ratio of short *MAC1* mRNA isoform to long isoform was 5.5 (± 2.3) in wild type strains and was 12.2 (± 2.2) in NMD mutants in CM. In low copper conditions, the short *MAC1* isoform accumulated 1.8 (± 1.1) fold higher than the long isoform in the wild-type strain, while the short *MAC1* isoform accumulated 3.0 (± 1.3) fold higher than the long isoform in NMD mutants (Supplementary Table 2).

To ensure that the presence of the longer *MAC1* mRNA isoform in these half-life measurements was not due to the *rpb1-1* yeast strains, we measured the half-life of *MAC1* mRNA in CM using wild-type (*UPF1*) and NMD mutant (*upf1Δ*) yeast strains treated with thiolutin. The W303a wild-type and NMD mutant strains lacking *rpb1-1* were used (Supplementary Table S1). The two strains were the same strains used to measure *MAC1* mRNA steady-state accumulation levels. The two *MAC1* mRNA isoforms were detected after thiolutin was used to terminate transcription in yeast cells. These two *MAC1* mRNA isoforms were similar to those observed with W303 *rpb1-1* strains grown in CM (Figure 2C and Supplementary Figure 1C). Specifically, the half-life of the major *MAC1* mRNA transcript in the wild-type strain was 10.1 (± 0.4) mins relative to 15.7 (± 1.2) mins in the NMD mutants (p-value = 0.007) (Figure 2C and Table 2). The half-lives measured using thiolutin were longer than those measured using *rpb1-1* to inhibit transcription (Figure 2A and 2C). These results are consistent with a previous study showing that thiolutin is also an inhibitor of mRNA degradation, and the apparent half-life measured in individual mRNA stability are probably longer than those measured using other techniques (30). These observations confirmed that isoforms of *MAC1* mRNA were not due to the yeast strains but possible due to copper deplete conditions, and that *MAC1* mRNA is a direct NMD target in CM.

Because *MAC1* mRNA does not contain introns, the generation of two *MAC1* mRNA isoforms was not due alternative splicing. We hypothesized that the two *MAC1* transcripts undergo alternative polyadenylation and vary in the length of their 3′-UTRs. 3′RACE analysis showed that, the *MAC1* transcript that was predominant in CM is 1.6 kb and has a 3’-UTR of ∼300 nt, while the 2.9 kb *MAC1* transcript had a 3′-UTR of ∼ 1400 nt (Figure 2D lane 1 and 2). The 3′RACE results showed that the ∼ 1400 nt PCR product was only detected in low copper conditions. There was an additional 3′RACE PCR product of ∼650 nt detected. This is most likely a non-specific band because in both CM and low copper conditions, a transcript of ∼ 1.9 kb, which would correspond to a 3′-UTR length of ∼ 650 nt, was not detected.

Additionally, the half-lives of the *MAC1* transcripts under high copper (600 μM Cu) were measured using identical time points as those used to measure the half-lives in CM and low copper (Figure 2E and Supplementary Figure S1D). The decay rate of the major *MAC1* mRNA isoforms surprisingly varied from the decay rate in CM. Notably, the longer *MAC1* isoform was not consistently detectable under high copper (600 μM Cu). Moreover, the longer *MAC1* mRNA is also undetectable in 600 μM copper steady-state northern blots. The half-life of the major *MAC1* mRNA isoform in wild-type strains was 30.8 (± 4.7) mins relative to 28.2 (± 6.1) mins in NMD mutants (Figure 2E and Table 2). These results corroborated the results of the steady-state accumulation levels showing that *MAC1* mRNA is only regulated by NMD under CM, but not in low copper or high copper conditions. In addition, *MAC1* mRNA is a direct NMD target under normal growth conditions (CM). Furthermore, the longer *MAC1* mRNA isoform is highly expressed only under low copper conditions, suggesting that the *MAC1* transcript undergoes alternative polyadenylation only under low copper.

### The *MAC1* 3′-UTR targets NMD insensitive *CYC1* mRNA to NMD-mediated degradation

Because *MAC1* mRNAs are differently regulated by NMD under various copper levels, we examined the functionality of the NMD-targeting features within the *MAC1* mRNAs. *MAC1* mRNAs have one potential NMD-targeting feature (Figure 1A). We investigated the role the atypical long 3′-UTRs plays in the NMD-mediated degradation of the *MAC1* mRNAs.

We hypothesized that the atypically long 3′-UTRs (300 and 1400nt, respectively) of the *MAC1* mRNAs contribute to the degradation of the mRNAs by NMD (Figure 1A). This hypothesis was based on our previous studies showing that atypically long 3′-UTR on other mRNAs involved in copper homeostasis affect their regulation by NMD (15,17). Furthermore, previous studies have demonstrated that replacement of the 3′-UTR of an NMD-insensitive mRNA with an atypical long 3′-UTR can cause the mRNA to be targeted by NMD (15,27,28). We generated two fusion mRNAs (Supplementary Figure 2). The *CYC1MAC1* 3′-UTR fusion mRNA was generated by replacing the 3′-UTR of *CYC1* with 1400 nt downstream of the *MAC1* STOP codon containing the *MAC1* 3′-UTR (Figure 3A). The *CYC1* mRNA, which encodes for iso-1-cytochrome C, was utilized because it has previously been used to study NMD cis-elements, and it is insensitive to the pathway (31,32).

**Figure 3.**
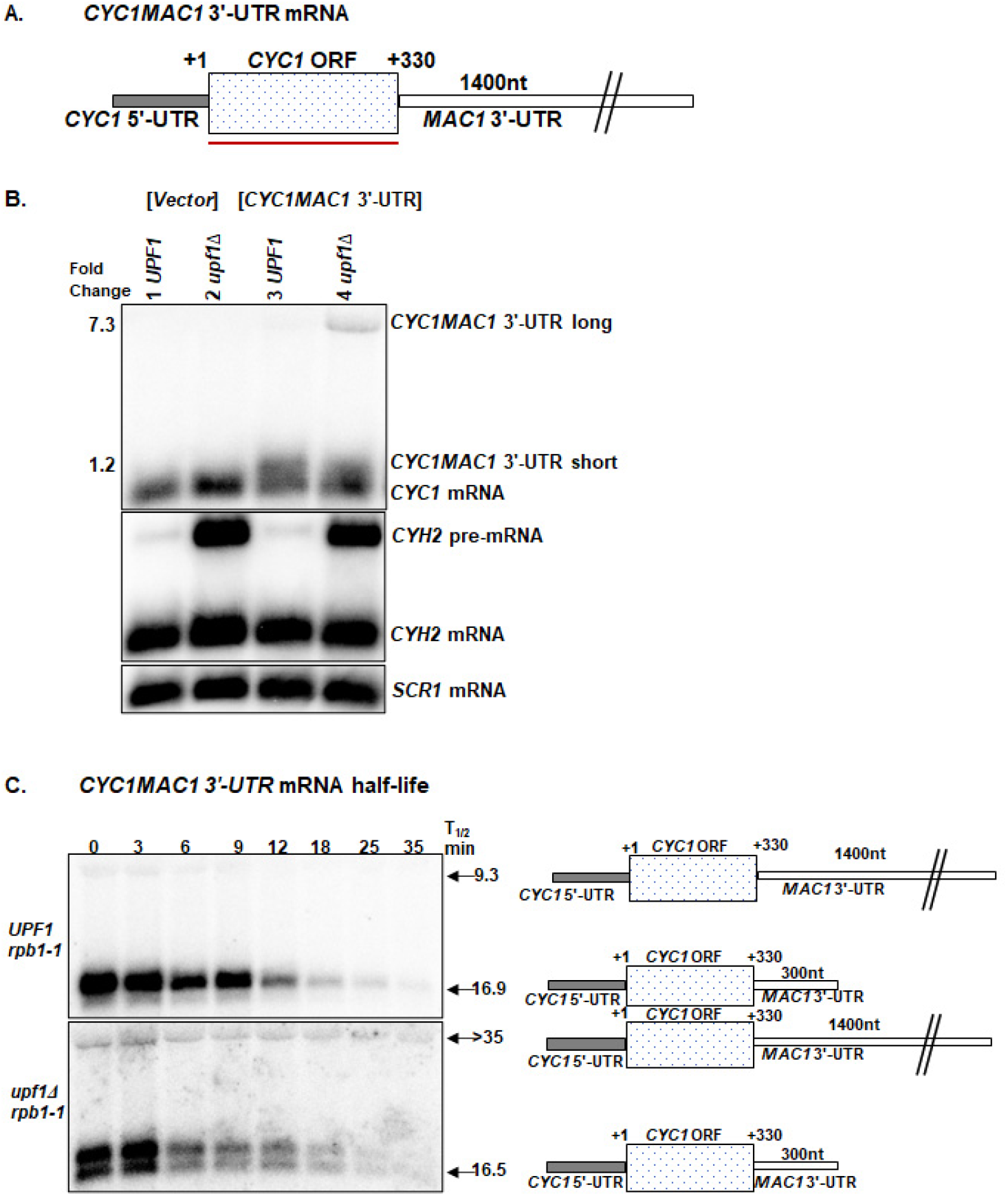
Schematic representation of *CYC1MAC1* 3′-UTR (A). Representative mRNA steady-state accumulation levels of *CYC1MAC1* 3′-UTR in CM -leu (B). The first two lanes of the steady-state northern blot (B) are loaded with RNA from yeast strains lacking the *CYC1MAC1* 3′-UTR construct and transformed with pRS315 (Vector control). The fold change (*upf1Δ/UPF1*) of *CYC1MAC1* 3′-UTR mRNA isoforms are shown left of the northern blot (B). Representative half-life northern blots of *CYC1MAC1* 3′-UTR mRNA in CM -leu (C). The northern blots were probed with DNA specific to the *CYC1* ORF indicated by red line. *CYH2* and *SCR1* were used as controls as described in Figure 1 (Supplementary Figure 3).

To find out if the *CYC1* mRNA was regulated by NMD due to the presence of the *MAC1* 3′-UTR, the levels of the *CYC1MAC1* 3′-UTR mRNA were measured in wild-type and NMD mutants. The *CYC1MAC1* 3′-UTR fusion construct encoded two mRNAs of ∼ 630 nt and ∼1730 nt, which is consistent with mRNAs containing a 3′-UTR of 300 nt and 1400 nt, respectively (Figure 3B, Supplementary Figure 2A and 3A), suggesting the alternative polyadenylation that *MAC1* undergoes is transferable to *CYC1*. The longer *CYC1MAC1* 3′-UTR mRNA isoform accumulated 7.3 (± 0.3) fold higher (p-value = 0.003) in NMD mutants relative to wild-type strains (Figure 3B and Table 3). Because the short *CYC1MAC1* 3′-UTR fusion isoform overlapped with *CYC1* mRNA, and *CYC1* mRNA is not regulated by NMD, we quantified the overlapped bands of *CYC1* mRNA and the short *CYC1MAC1* 3′-UTR isoform to calculate the fold change. Unlike *MAC1* mRNAs, the short *CYC1MAC1* 3′-UTR mRNA did not accumulate to significantly higher levels in NMD mutants. This *CYC1MAC1* 3′-UTR fusion mRNA accumulated 1.2 (± 0.2) fold higher in NMD mutants relative to the wild-type strain (Figure 3B and Table 3).

**Table 3.** mRNA steady-state accumulations and half-lives of the *CYC1MAC1 3′-UTR* and *MAC1CYC1 3′-UTR* mRNAs. Wild-type (W303) and NMD mutants (AAY320) were used to measure mRNA steady-state accumulation levels, while AAY334 and AAY335 transformed with the respective constructs were used to determine half-lives. All yeast strains used were grown under regular conditions lacking leucine in complete minimal (CM) media and 100 μM BCS (low Cu). The steady-state and half-life experiments were done in triplicate and are reported as averages ± standard deviation. ND, not determined

| Growth media | Name | Relative mRNA accumulation<br>(*upf1Δ*/*UPF1*) | mRNA half-life (min)<br>*UPF1* | mRNA half-life (min)<br>*upf1Δ* |
| --- | --- | --- | --- | --- |
| **CM** | *CYC1MAC1 3'-UTR* long | 7.3 (± 0.3) | 9.3 (± 3.3) | > 35 |
| | *CYC1MAC1 3'-UTR* short | 1.2 (± 0.2) | 16.9 (± 1.7) | 16.5 (± 3.3) |
| | *MAC1CYC1 3'-UTR* | 0.6 (± 0.05) | 5.0 (± 0.5) | 3.7 (± 1.0) |
| **Low Cu** | *CYC1MAC1 3'-UTR* long | 8.8 (± 4.7) | ND | > 35 |
| | *CYC1MAC1 3'-UTR* short | 0.9 (± 0.3) | 17.4 (± 4.1) | 19.4 (± 8.5) |
| | *MAC1CYC1 3'-UTR* | 1.3 (± 0.4) | 33.3 (± 13.7) | 15.4 (± 1.4) |

Additionally, the long *CYC1MAC1* 3′-UTR isoform was a directly regulated by NMD with significantly different decay rates in wild-type and NMD mutant strains (Figure 3C and Supplementary Figure 3B). The half-life of the long *CYC1MAC1* 3′-UTR mRNA in the wild-type strain was 9.3 (± 3.3) mins, while in the NMD mutants the half-life of the long *CYC1MAC1* 3′-UTR mRNA transcript could not be determined because it did not decay within the 35-min period (Table 3). No significant difference in decay rate of the short *CYC1MAC1* 3′-UTR was detected between wild-type (16.9 (± 1.7) mins) and NMD mutant strains (16.5 (± 2.5) mins) (Figure 3C and Supplementary Figure 3B).

### Heterologous *MAC1* 3′-UTR expression is not responsive to copper deficiency

Under low copper conditions, the endogenous *MAC1* transcript undergoes alternative 3′-end processing generating two major mRNA isoforms. Importantly, both *MAC1* mRNA isoforms escape NMD-mediated regulation in these conditions (Figure 2B). To determine whether heterologous *MAC1* 3′-UTR in *CYC1* mRNA acted similarly, the levels of the *CYC1MAC1* 3′-UTR fusion mRNA were measured under low copper conditions. Because the endogenous *MAC1* mRNAs were stabilized under low copper conditions, we hypothesized that the 3′-UTRs of *MAC1* mRNA could contribute to the stabilization of *MAC1* mRNAs.

Under low copper conditions, the *CYC1MAC1* 3′-UTR fusion construct encoded two mRNA isoforms similar to CM. The long *CYC1MAC1* 3′-UTR fusion mRNA accumulated 8.8 (± 4.7) fold in NMD mutants relative to wild type strains (Figure 4A). The short *CYC1MAC1* 3′-UTR mRNA did not accumulate to higher levels in NMD mutants relative to wild type in low copper conditions. This is the same as what we observed in CM (Figure 3A and 4A). In addition, the endogenous *CYC1* was downregulated under low copper conditions relative to CM (Figure 4A and Supplementary Figure 4A). Furthermore, no significant difference in half-life of the short *CYC1MAC1* 3′-UTR fusion mRNA was observed in low copper conditions. The short *CYC1MAC1* 3′-UTR was not detected to have significantly different decay rate relative to CM, nor between wild-type and NMD mutant strains under low copper conditions (Figure 3C and 4B). The long *CYC1MAC1* 3′-UTR isoform was a direct NMD target with altered decay rates in the wild-type and NMD mutant strain in low copper conditions (Figure 4B and Supplementary Figure 4B). However, the half-life of the long *CYC1MAC1* 3’-UTR mRNA in wild type strains was not determined because the relative low abundance of the long *CYC1MAC1* 3’-UTR mRNA. In the NMD mutant, the half-life of the long *CYC1MAC1* 3’-UTR mRNA was more than 35 mins. These results suggested that the *MAC1* 3′-UTR in the context of the *CYC1* mRNA is sensitive to NMD but does not respond to low copper conditions, like the endogenous *MAC1* mRNA.

**Figure 4.**
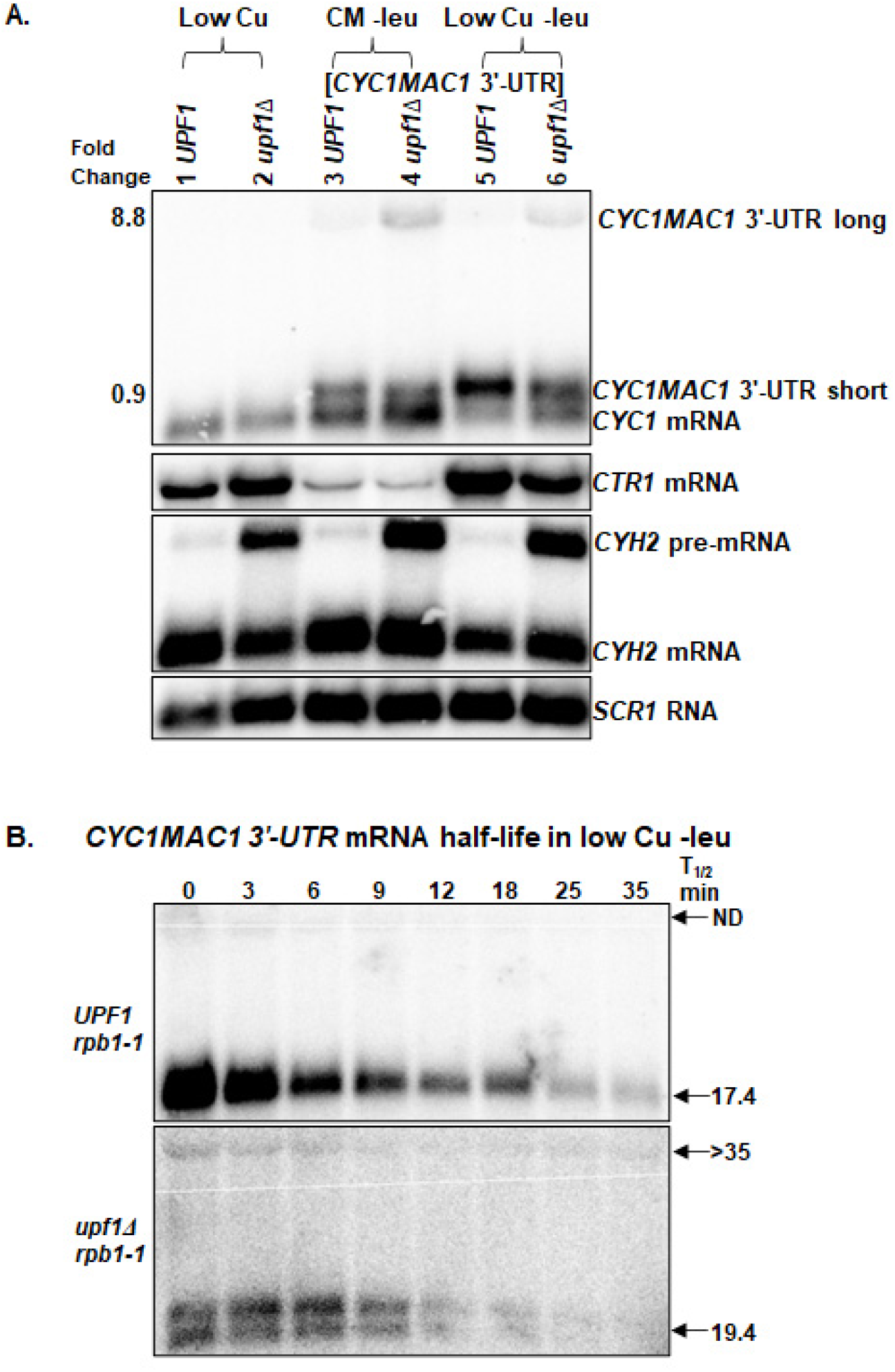
Representative mRNA steady-state accumulation levels of *CYC1MAC1* 3′-UTR mRNA in low copper (A). mRNA steady-state levels were measured with total RNA from wild-type strain W303 (*UPF1*) (39), and NMD mutants AAY320 (*upf1Δ*)(25). The first two lanes of the steady-state northern blots (A) are loaded with RNA from yeast strains lacking the *CYC1MAC1* 3′-UTR construct grown in low copper conditions. Lanes 3 and 4 are loaded with RNA from yeast strains with *CYC1MAC1* 3′-UTR construct grown in CM -leu. The fold change (*upf1Δ/UPF1*) of *CYC1MAC1* 3′-UTR mRNA isoforms under low copper conditions are shown left of the northern blot (A). Representative half-life northern blots of *CYC1MAC1* 3′-UTR mRNA in low copper (B). The northern blot was probed with DNA specific to *CYC1* ORFs. *CTR1, CYH2* and *SCR1* were used as controls as described in Figure 1 (Supplementary Figure 4).

### Substitution of the *MAC1* 3′-UTR with the *CYC1* 3′-UTR renders the *MAC1* mRNA insensitive to NMD

Subsequently, we investigated whether the *MAC1* mRNA could be regulated by NMD in the absence of the long 3′-UTRs. We expected that *MAC1* mRNAs would no longer be regulated by NMD because the mRNAs do not have additional recognizable NMD-targeting features (Figure 1A). To test this hypothesis, we generated the *MAC1CYC1* 3′-UTR fusion construct, which contains the 5′-UTR and ORF of *MAC1* fused to *CYC1* 3′-UTR (Figure 5A and Supplementary Figure 2B).

**Figure 5.**
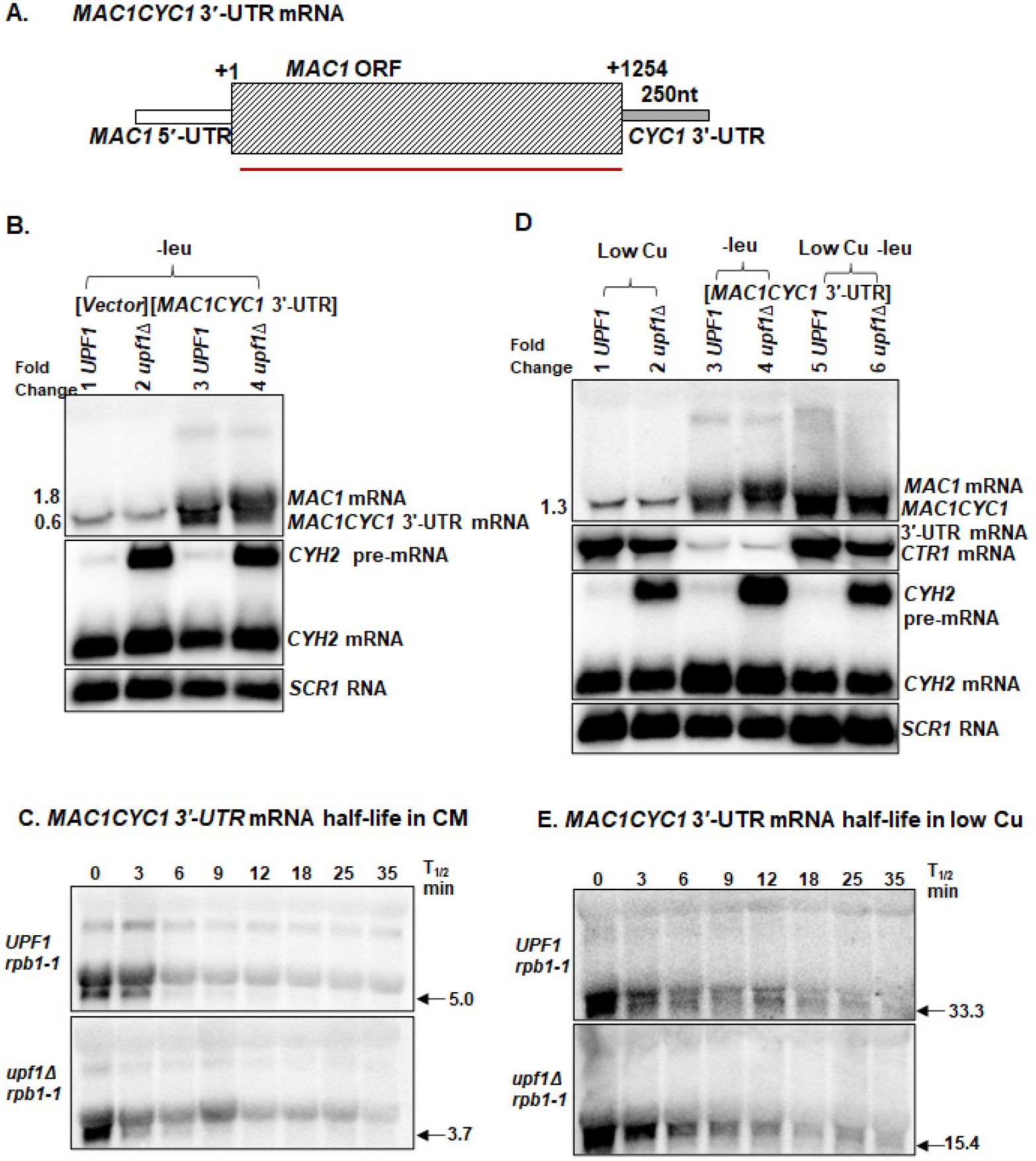
Schematic representation of *MAC1CYC1* 3′-UTR (A). Representative mRNA steady-state accumulation levels of *MAC1CYC1* 3′-UTR in CM -leu (B) and low copper -leu (D). The first two lanes of the steady-state northern blot (B) are loaded with RNA from a yeast strain lacking the *MAC1CYC1* 3′-UTR construct and transformed with pRS315 (Vector control). The *MAC1CYC1* 3′-UTR mRNA isoform fold change (*upf1Δ/UPF1*) are shown left of the northern blot (B). The first two lanes of the steady-state northern blot (D) are loaded with RNA from yeast strains lacking the *MAC1CYC1* 3′-UTR construct. Lanes 3 and 4 are loaded with RNA from yeast strains containing *MAC1CYC1* 3′-UTR grown in CM -leu. Representative half-life northern blots of *MAC1CYC1* 3′-UTR mRNA in CM -leu (C) and low copper -leu (E). The arrow indicates the half-life of mRNAs produced by *MAC1CYC1* 3′-UTR construct. The northern blots were probed with DNA specific to the *MAC1* ORF indicated by red line. *CTR1, CYH2*, and *SCR1* were used as controls as described in Figure 1 (Supplementary Figure 5).

The *MAC1CYC1* 3′-UTR fusion construct generated one transcript, unlike the endogenous *MAC1* which generates two low abundance transcripts in wild-type yeast strains. Furthermore, the *MAC1CYC1* 3′-UTR mRNA did not accumulate to significantly higher levels in NMD mutants (Figure 5B and Table 3). The steady-state accumulation levels of the endogenous *MAC1* mRNA isoform in wild-type and NMD mutants expressing the *MAC1CYC1* 3′-UTR fusion mRNA was 1.8 (± 0.3), which was not significantly different from the fold change we measured from yeast strains without the *MAC1CYC1 3’-UTR* mRNA (Figure 1B and Table 1), indicating that the *MAC1CYC1* 3′-UTR mRNA did not influence the regulation of endogenous *MAC1* mRNA. No significant difference in the half-life of *MAC1CYC1* 3′-UTR mRNA was observed (Figure 5C and Supplementary Figure 5A). *MAC1CYC1* 3′-UTR mRNA half -life was 5.0 (± 0.5) mins in wild-type strains relative to 3.7 (± 1.0) mins in NMD mutants (Figure 5C and Table 3). These results demonstrate that the *MAC1CYC1* 3′-UTR mRNA escapes NMD under these conditions.

### Copper deficiency stabilizes *MAC1* mRNA in the absence of the 3′-UTR

Because the endogenous *MAC1* mRNAs accumulated to higher levels in low copper conditions relative to CM, and the heterologous *MAC1* 3′-UTR in *CYC1* mRNA did not respond to low copper conditions, we were interested in examining how *MAC1* mRNA behaves under low copper conditions in the absence of the *MAC1* 3′-UTR. Wild-type (*UPF1*) and NMD mutants (*upf1*Δ) yeast strains containing plasmid-encoded *MAC1CYC1* 3′-UTR fusion construct were grown in complete minimal containing 100 mM BCS (low copper) lacking leucine.

*MAC1CYC1* 3′-UTR mRNA accumulated to higher levels in low copper conditions relative to CM (Figure 5D). However, the *MAC1CYC1* 3′-UTR mRNA did not accumulate to significantly higher levels in NMD mutants relative to the wild-type strains (Figure 5D lane 5 and 6). Furthermore, the endogenous *MAC1* mRNA did not accumulate to higher levels in NMD mutants relative to wild type in low copper conditions (Figure 1B lanes 3 and 4, Figure 5D lane 5 and 6). *CTR1* mRNA was used as the low copper control, as Mac1p activates *CTR1* mRNA under low copper conditions. The expression of *CTR1* mRNA in yeast strains containing the *MAC1CYC1* 3′-UTR fusion construct was comparable to strains lacking the *MAC1CYC1* 3′-UTR construct in low copper conditions (Figure 5D compare lanes 1 and 2 to lanes 5 and 6). Moreover, the mRNA encoded by *MAC1CYC1* 3′-UTR construct was stabilized in low copper conditions in both wild-type and NMD mutants relative to CM (Figure 5C, E and Supplementary Figure 5). Specifically, the half-life of *MAC1CYC1* 3′-UTR mRNA was 33.3 (± 13.7) mins in wild-type strains relative to 15.4 (± 1.4) mins in NMD mutants (Figure 5E and Table 3).

## DISCUSSION

The control of gene expression by NMD can be condition specific depending on the function of the regulated genes. Here we show that the regulation of the *MAC1* mRNAs by NMD in varying levels of copper is unique and complex (Figure 6). The short *MAC1* transcript is directly regulated by NMD in regular growth conditions, however it escapes NMD-mediated degradation in low and high copper. Copper levels affect the expression of *MAC1* mRNA isoforms and sensitivity to NMD. The long *MAC1* transcript is of low abundance in complete minimal and high copper, while the expression level of this isoform is elevated under low copper conditions. Notably, the short *MAC1* transcript is not an NMD target but is stabilized under low and high copper conditions (Figure 2B and 2E). Because Mac1p is a transcription factor that activates genes under low copper conditions we would expect the downstream genes to be regulated by NMD like *MAC1*, although they would be indirect NMD targets. *CTR1* and *FRE1* mRNAs are well characterized Mac1p targets and they should be indirect NMD targets under normal growth conditions where *MAC1* mRNA is directly regulated by the pathway according to prototypical direct and indirect NMD targets (Figure 1B (16)). However, the model for direct and indirect NMD targets is not applicable here because the functionality of Mac1p is dependent on copper levels.

**Figure 6.**
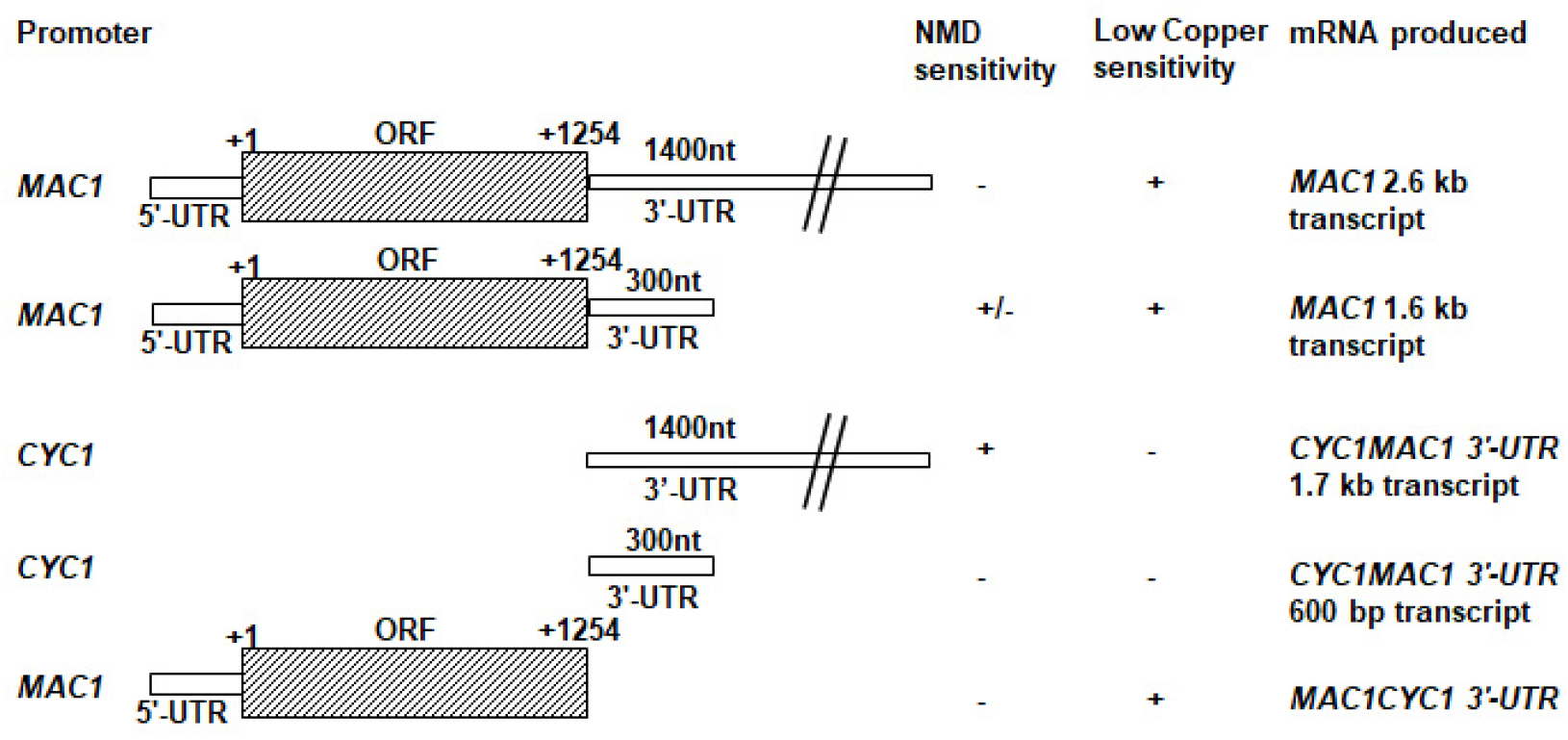
Schematic summary of the *MAC1* mRNA analysis. A schematic representation of the fragments of the *MAC1* mRNAs that are found in each of the fusion mRNAs, the promoter driving the expression of each mRNA, and the mRNA that lead to the inference on the level of NMD and copper sensitivity is shown. NMD sensitivity of + indicates that the mRNA containing that fragment of *MAC1* was sensitive to NMD. +/− indicates that the fragment of *MAC1* was sensitive to NMD in normal growth condition but evades NMD in low copper and high copper conditions. – indicates that the mRNA was insensitive to the NMD pathway. Copper sensitivity of + indicates that the mRNA containing that fragment of *MAC1* was responsive to low copper condition. – indicates that the mRNA was not responsive to low copper condition. The numbers on the schematic images are based on the *MAC1* ORF start point at +1.

There are differing reports on the effect of copper on Mac1p. A previous study reported that copper differently regulates the degradation of Mac1p (33). The same study found that the expression of a plasmid encoded *MAC1* mRNA did not change with increase in copper levels, but Mac1p was degraded in a copper ion concentration-dependent manner (33). A separate study reported that Mac1p levels increased in copper replete conditions (34). Furthermore, Mac1p is inactive when bound to Cu(I) and may not be functional under high copper conditions (21). Our study found that elevated copper levels do not inhibit *MAC1* mRNA expression, however elevated copper levels inhibit Mac1p activation of *CTR1* (Figure 1B). Thus, it appears that in elevated copper levels, *MAC1* mRNA is transcribed and translated, but Mac1p is not functional in order to inhibit copper induced toxicity due to continuous copper uptake by transporter activity (33). Furthermore, under elevated copper levels, metallothioneins such as *CUP1* and *CRS5*, and additional genes involved in adaptation to high copper levels are elevated (Figure 1B). We have reported previously that wild-type and NMD mutant yeast strains tolerate high copper levels with the NMD mutants being more tolerant than the wild-type strains (35).

Additionally, heterologous expression of the *MAC1* 3′-UTR on *CYC1* showed that under normal growth conditions the alternative 3′-end processing that *MAC1* undergoes is transferable. We expected that the *MAC1* 3′-UTR would be sufficient to target an NMD-insensitive mRNA to pathway. However, the 1400 nt *MAC1* 3′-UTR was sufficient to target an NMD-insensitive mRNA to the pathway, while the 300 nt *MAC1* 3′-UTR did not target *CYC1* mRNA to the pathway. Conversely, under low copper conditions, both *MAC1* mRNA isoforms escape NMD-mediated degradation. Interestingly, the long *CYC1MAC*1 3′-UTR transcript was regulated by NMD under these conditions suggesting that escape from NMD requires the *MAC1* 5′-UTR and ORF. Furthermore, in the context of *MAC1* the 300 nt 3’-UTR was sufficient to target the mRNA to NMD under normal growth conditions but in the context of *CYC1* this 3’-UTR was inadequate to target the mRNA to NMD under the conditions tested here.

We also observed that under low copper the endogenous *CYC1* mRNA accumulates to lower levels relative to normal copper levels (Figure 4A and Supplementary Figure 4A). *CYC1* encodes for one of the key components of the mitochondrial respiratory chain. A previous study reported that *CYC1* mRNA was downregulated under copper deprivation relative to CM, which is consistent with what we observed here (Figure 4A and Supplementary Figure 4A) (36). However, the levels of the *CYC1MAC1* 3′-UTR mRNA were elevated under low copper in both the wild-type and NMD mutant although only the longer *CYC1MAC1* 3′-UTR was regulated by NMD (Figure 4A)

The absence of the *MAC1* 3′-UTR makes the mRNA insensitive to NMD, however it is responsive to low copper conditions. We observed that the *MAC1CYC1* 3′-UTR mRNA does not accumulate to higher levels in NMD mutants relative to wild-type strains. Moreover, the half-life of *MAC1CYC1* 3′-UTR mRNA was shorter in the NMD mutant relative to the wild-type strain in normal growth conditions (Figure 5C). These observations suggest that the *CYC1* 3′-UTR could have a destabilizing effect on the *MAC1* transcript, although that destabilization was independent of Upf1p and therefore unrelated to NMD. Comparable findings have been reported with the 3′-UTR from *CYC1* mRNA accelerating the normal mechanism of mRNA decay (31).

*MAC1* mRNA lacking the 3′-UTR responds to the low copper conditions due to transcription and stabilization of the mRNA. That is, the steady-state accumulation levels of *MAC1* mRNA increase due to increased transcription in low copper conditions. Subsequently, *MAC1* mRNA is stabilized and translated. The produced Mac1p activates the transcription of *CTR1*. This conclusion is based on the observation that the expression of *CTR1* mRNA accumulated ∼10-fold higher in strains containing plasmid-encoded *MAC1CYC1* 3′-UTR fusion mRNA in low copper conditions relative to normal growth conditions (Figure 5D and Supplementary Table 3). As stated above, *CTR1* encodes a required for high-affinity copper uptake and is activated by Mac1p. A previous study reported that overexpression of *MAC1* increased *CTR1* expression level in low copper conditions (20). Additionally, these observations suggest that because *MAC1CYC1* 3′-UTR mRNA is most likely efficiently translated in low copper conditions, this protects the mRNA from NMD-mediated degradation. Previous studies found that the normal-looking mRNA that are less efficiently translated are regulated by NMD. Conversely, well translated NMD targets may escape NMD based on the conditions (37). Furthermore, the half-life of *MAC1CYC1* 3′-UTR mRNA retains the response to low copper conditions like the endogenous *MAC1* mRNA (Figure 2B and 5E). The decay rate of the *MAC1CYC1* 3′-UTR mRNA is much longer under low copper relative to CM, suggesting that the mRNA is stabilized in low copper relative to normal growth conditions (Figure 5C and 6B).

This study demonstrates that regulation of genes involved in bio-metal homeostasis occurs at multiple levels. Specifically, precise regulation of *MAC1* in response to changing copper levels occurs at transcription, mRNA stability and protein levels. *MAC1* mRNA is stabilized and efficiently translated under low copper conditions. Efficient translation of *MAC1* mRNA under low copper conditions could protect the mRNA from NMD. Mac1p activates *CTR1* in response to low copper conditions to uptake copper from the environment. The deletion of *MAC1* results in the absence of *CTR1* mRNA (38).

The sensitivity of *MAC1* to NMD is due to the 3′-UTR, while sensitivity to copper is at transcriptional and posttranslational levels. *MAC1* mRNA lacking the 3′-UTR is no longer regulated by NMD in the conditions tested here. Altogether, *MAC1* 3′-UTR targets the mRNA to NMD under normal growth conditions (Figure 7A). When yeast cells are grown under low copper and high copper conditions, *MAC1* mRNA is likely efficiently translated, though Mac1p is non-functional in elevated copper levels and does not activate the expression of *CTR1* and *FRE1* (33) (Figure 7B and 7C).

**Figure 7.**
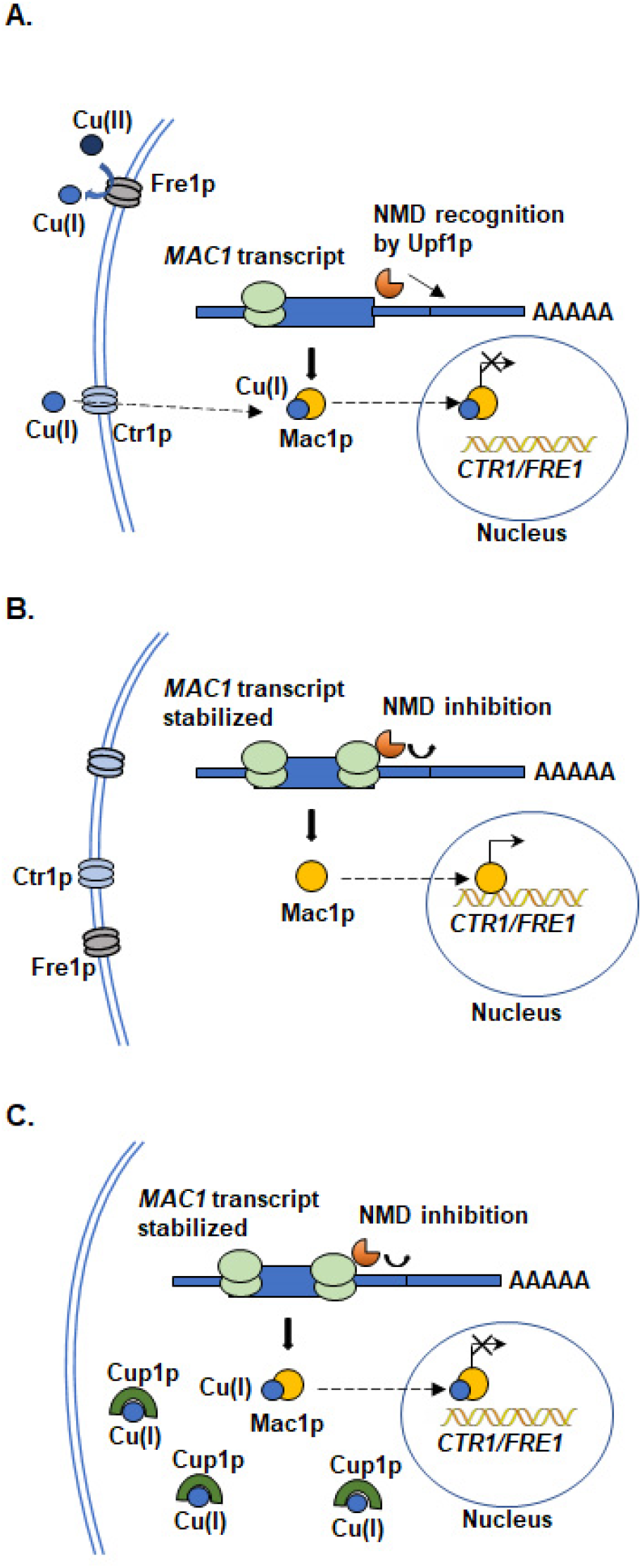
A proposed model to explain the mechanisms of why *MAC1* mRNA is stabilized under specific conditions. Schematic representation of *MAC1* mRNA being degraded by NMD in CM (A). *MAC1* mRNA is produced and transported from nucleus to the cytoplasm. Subsequently, *MAC1* mRNA is translated to generate Mac1 protein. Mac1p binding to Cu(I) is transported into nucleus but no longer activates the transcription of *CTR1* and *FRE1*, thus maintaining the appropriate intracellular copper levels. Schematic representation of *MAC1* mRNA which is resistant to NMD in low copper conditions (B). *MAC1* mRNA is efficiently transcribed and translated under low copper conditions. Efficient translation of *MAC1* mRNA may possibly protect the mRNA from degradation by NMD. The Mac1p is imported into the nucleus and activates the transcription of *CTR1* and *FRE1* to uptake the environmental copper. Schematic representation of *MAC1* mRNA evading NMD in high copper conditions (C). Under high copper conditions, *MAC1* mRNA is transcribed and translated. Elevated copper might influence *MAC1* translation and protect the mRNA from NMD. Although Mac1p is produced, Mac1p does not activate the transcription of *CTR1* and *FRE1*. In this conditions, Cup1p metallothionein chelates extra Cu(I) to prevent copper induced toxicity of yeast cells.

Translation of specific mRNAs with NMD targeting features may be altered in response to changing environmental copper levels resulting in protection of mRNAs from rapid decay by the NMD pathway. This example from the yeast transcriptome demonstrating that copper levels likely influence *MAC1* mRNA processing and expression levels, thus influencing NMD-mediated degradation may also occur with other mRNAs involved in metal ion homeostasis.

## SUPPLEMENTARY DATA

Supplementary Data are available at NAR online.

## ACKNOWLEDGEMENT

We are grateful to Ethan Blasdel for critically reading the manuscript. We also thank the Molecular Biosciences Center (MBC) facility at Baylor University for equipment and supplies.

## REFERENCES

1. Maquat, L.E. (1995) When cells stop making sense: effects of nonsense codons on RNA metabolism in vertebrate cells. RNA, 1, 453–465.

2. He, F., Peltz, S.W., Donahue, J.L., Rosbash, M. and Jacobson, A. (1993) Stabilization and ribosome association of unspliced pre-mRNAs in a yeast upf1-mutant. Proc Natl Acad Sci U S A, 90, 7034–7038.

3. He, F., Li, X., Spatrick, P., Casillo, R., Dong, S. and Jacobson, A. (2003) Genome-wide analysis of mRNAs regulated by the nonsense-mediated and 5’ to 3’ mRNA decay pathways in yeast. Mol Cell, 12, 1439–1452.

4. Mitrovich, Q.M. and Anderson, P. (2000) Unproductively spliced ribosomal protein mRNAs are natural targets of mRNA surveillance in C. elegans. Genes Dev, 14, 2173–2184.

5. Thompson, D.M. and Parker, R. (2007) Cytoplasmic decay of intergenic transcripts in Saccharomyces cerevisiae. Mol Cell Biol, 27, 92–101.

6. Wang, J., Vock, V.M., Li, S., Olivas, O.R. and Wilkinson, M.F. (2002) A quality control pathway that down-regulates aberrant T-cell receptor (TCR) transcripts by a mechanism requiring UPF2 and translation. J Biol Chem, 277, 18489–18493.

7. Welch, E.M. and Jacobson, A. (1999) An internal open reading frame triggers nonsense-mediated decay of the yeast SPT10 mRNA. Embo j, 18, 6134–6145.

8. Celik, A., He, F. and Jacobson, A. (2017) NMD monitors translational fidelity 24/7. Curr Genet, 63, 1007–1010.

9. Mendell, J.T., Sharifi, N.A., Meyers, J.L., Martinez-Murillo, F. and Dietz, H.C. (2004) Nonsense surveillance regulates expression of diverse classes of mammalian transcripts and mutes genomic noise. Nat Genet, 36, 1073–1078.

10. Guan, Q., Zheng, W., Tang, S., Liu, X., Zinkel, R.A., Tsui, K.W., Yandell, B.S. and Culbertson, M.R. (2006) Impact of nonsense-mediated mRNA decay on the global expression profile of budding yeast. PLoS Genet, 2, e203.

11. Rehwinkel, J., Letunic, I., Raes, J., Bork, P. and Izaurralde, E. (2005) Nonsense-mediated mRNA decay factors act in concert to regulate common mRNA targets. RNA, 11, 1530–1544.

12. Johansson, M.J., He, F., Spatrick, P., Li, C. and Jacobson, A. (2007) Association of yeast Upf1p with direct substrates of the NMD pathway. Proc Natl Acad Sci U S A, 104, 20872–20877.

13. He, F. and Jacobson, A. (1995) Identification of a novel component of the nonsense-mediated mRNA decay pathway by use of an interacting protein screen. Genes Dev, 9, 437–454.

14. Cui, Y., Hagan, K.W., Zhang, S. and Peltz, S.W. (1995) Identification and characterization of genes that are required for the accelerated degradation of mRNAs containing a premature translational termination codon. Genes Dev, 9, 423–436.

15. Peccarelli, M., Scott, T.D., Steele, M. and Kebaara, B.W. (2016) mRNAs involved in copper homeostasis are regulated by the nonsense-mediated mRNA decay pathway depending on environmental conditions. Fungal Genet Biol, 86, 81–90.

16. Peccarelli, M., Scott, T.D. and Kebaara, B.W. (2019) Nonsense-mediated mRNA decay of the ferric and cupric reductase mRNAs FRE1 and FRE2 in Saccharomyces cerevisiae. FEBS Lett, 593, 3228–3238.

17. Peccarelli, M., Scott, T.D., Wong, H., Wang, X. and Kebaara, B.W. (2014) Regulation of CTR2 mRNA by the nonsense-mediated mRNA decay pathway. Biochim Biophys Acta, 1839, 1283–1294.

18. Deliz-Aguirre, R., Atkin, A.L. and Kebaara, B.W. (2011) Copper tolerance of Saccharomyces cerevisiae nonsense-mediated mRNA decay mutants. Curr Genet, 57, 421–430.

19. De Freitas, J., Wintz, H., Kim, J.H., Poynton, H., Fox, T. and Vulpe, C. (2003) Yeast, a model organism for iron and copper metabolism studies. Biometals, 16, 185–197.

20. Yamaguchi-Iwai, Y., Serpe, M., Haile, D., Yang, W., Kosman, D.J., Klausner, R.D. and Dancis, A. (1997) Homeostatic regulation of copper uptake in yeast via direct binding of MAC1 protein to upstream regulatory sequences of FRE1 and CTR1. J Biol Chem, 272, 17711–17718.

21. Jensen, L.T. and Winge, D.R. (1998) Identification of a copper-induced intramolecular interaction in the transcription factor Mac1 from Saccharomyces cerevisiae. EMBO J, 17, 5400–5408.

22. Peña, M.M., Koch, K.A. and Thiele, D.J. (1998) Dynamic regulation of copper uptake and detoxification genes in Saccharomyces cerevisiae. Mol Cell Biol, 18, 2514–2523.

23. Martins, L.J., Jensen, L.T., Simon, J.R., Keller, G.L. and Winge, D.R. (1998) Metalloregulation of FRE1 and FRE2 homologs in Saccharomyces cerevisiae. J Biol Chem, 273, 23716–23721.

24. Wong, A., Lam, E.M., Pai, C., Gunderson, A., Carter, T.E. and Kebaara, B.W. (2021) Variation of the response to metal ions and nonsense-mediated mRNA decay across different Saccharomyces cerevisiae genetic backgrounds. Yeast, 38, 507–520.

25. Kebaara, B., Nazarenus, T., Taylor, R. and Atkin, A.L. (2003) Genetic background affects relative nonsense mRNA accumulation in wild-type and upf mutant yeast strains. Curr Genet, 43, 171–177.

26. Peccarelli, M. and Kebaara, B.W. (2014) Measurement of mRNA decay rates in Saccharomyces cerevisiae using rpb1-1 strains. J Vis Exp.

27. Murtha, K., Hwang, M., Peccarelli, M.C., Scott, T.D. and Kebaara, B.W. (2019) The nonsense-mediated mRNA decay (NMD) pathway differentially regulates COX17, COX19 and COX23 mRNAs. Curr Genet, 65, 507–521.

28. Kebaara, B.W. and Atkin, A.L. (2009) Long 3’-UTRs target wild-type mRNAs for nonsense-mediated mRNA decay in Saccharomyces cerevisiae. Nucleic Acids Res, 37, 2771–2778.

29. Sikorski, R.S. and Hieter, P. (1989) A system of shuttle vectors and yeast host strains designed for efficient manipulation of DNA in Saccharomyces cerevisiae. Genetics, 122, 19–27.

30. Pelechano, V. and Perez-Ortin, J.E. (2008) The transcriptional inhibitor thiolutin blocks mRNA degradation in yeast. Yeast, 25, 85–92.

31. Muhlrad, D. and Parker, R. (1999) Aberrant mRNAs with extended 3’ UTRs are substrates for rapid degradation by mRNA surveillance. RNA, 5, 1299–1307.

32. Zaret, K.S. and Sherman, F. (1984) Mutationally altered 3’ ends of yeast CYC1 mRNA affect transcript stability and translational efficiency. J Mol Biol, 177, 107–135.

33. Zhu, Z., Labbe, S., Pena, M.M. and Thiele, D.J. (1998) Copper differentially regulates the activity and degradation of yeast Mac1 transcription factor. J Biol Chem, 273, 1277–1280.

34. Keller, G., Bird, A. and Winge, D.R. (2005) Independent metalloregulation of Ace1 and Mac1 in Saccharomyces cerevisiae. Eukaryot Cell, 4, 1863–1871.

35. Wong, A., Lam, E.M., Pai, C., Gunderson, A., Carter, T.E. and Kebaara, B.W. (2021) Variation of the response to metal ions and nonsense-mediated mRNA decay across different Saccharomyces cerevisiae genetic backgrounds. Yeast.

36. van Bakel, H., Strengman, E., Wijmenga, C. and Holstege, F.C. (2005) Gene expression profiling and phenotype analyses of S. cerevisiae in response to changing copper reveals six genes with new roles in copper and iron metabolism. Physiol Genomics, 22, 356–367.

37. Celik, A., Baker, R., He, F. and Jacobson, A. (2017) High-resolution profiling of NMD targets in yeast reveals translational fidelity as a basis for substrate selection. RNA, 23, 735–748.

38. Wood, L.K. and Thiele, D.J. (2009) Transcriptional activation in yeast in response to copper deficiency involves copper-zinc superoxide dismutase. J Biol Chem, 284, 404–413.

39. Wente, S.R., Rout, M.P. and Blobel, G. (1992) A new family of yeast nuclear pore complex proteins. J Cell Biol, 119, 705–723.

