## Supplementary information for "Nonsense-mediated mRNA decay of metal-binding activator *MAC1* is dependent on copper levels and 3′-UTR length in *Saccharomyces cerevisiae*"

**Supplementary Table 1.** *Saccharomyces cerevisiae* strains used in this study.

| Yeast Strain | Genotype | Source |
|---|---|---|
| **W303** | *a, ade2-1, ura3-1, his3-11,15, trp1-1, leu2-3,112, can1-101* | Wente *et al*. 1992 |
| **AAY320** | *a, ade2-1, ura3-1, his3-11,15, trp1-1, leu2-3,112, can1-100, UPF1::URA3 (upf1-Δ2)* | Kebaara *et al*., 2003 |
| **AAY334** | *a, ADE2, ura3-1 or ura3-52, his3-52, his3-11,15; trp1-1, leu2-3,112, rpb1-1* | Kebaara *et al*., 2003 |
| **AAY335** | *a, ADE2, ura3-1 or ura3-52, his3-52, his3-11,15; trp1-1, leu2-3,112, rpb1-1, upf1-Δ2 (URA3)* | Kebaara *et al*., 2003 |

**Supplementary Table 2.** Relative mRNA accumulation levels of short *MAC1* isoform versus long *MAC1* isoform at 0 time point in half-life measurements. Wild-type (AAY334) and NMD mutants (AAY335) have the *rpb1-1* allele. All yeast strains used were grown under standard growth conditions in complete minimal media with 100 µM BCS (low Cu). The ratios of *MAC1* short isoform to *MAC1* long isoform were done in triplicate at 0 time point of half-life experiments and are reported as averages ± standard deviation.

| Growth condition | AAY334 (*UPF1*) (Short *MAC1*/Long *MAC1*) | AAY335 (*upf1Δ*) (Short *MAC1*/Long *MAC1*) |
|---|---|---|
| **CM** | 5.5 (± 2.3) | 12.2 (± 2.2) |
| **100 µM BCS*** | 1.8 (± 1.1) | 3.0 (± 1.3) |

**Supplementary Table 3.** Relative mRNA accumulation levels of *CTR1* from Figure 5D. Wild-type (W303) and NMD mutants (AAY320) were used to determine mRNA steady-state accumulation levels. Yeast strains transformed with *MAC1CYC1* 3′-UTR were grown under regular growth conditions lacking leucine in complete minimal (CM -leu) media and CM -leu containing 100 µM BCS (low Cu -leu). Yeast strains without the *MAC1CYC1* 3′-UTR construct were grown under regular growth conditions containing 100 µM BCS (low Cu). Lane 1 and 2 of the steady-state northern blots (Figure 5D) are loaded with RNA from wild-type and *upf1Δ* yeast strains lacking the *MAC1CYC1* 3′-UTR construct grown in low copper conditions. Lanes 3 and 4 are loaded with RNA from wild-type and *upf1Δ* yeast strains with *MAC1CYC1* 3′-UTR construct grown in CM -leu. Lane 5 and 6 are loaded with RNA from wild-type and *upf1Δ* yeast strains with *MAC1CYC1* 3′-UTR construct grown in low Cu -leu. The steady-state accumulation levels of *MAC1CYC1* 3′-UTR construct grown in low Cu -leu were done in triplicate.

| **Yeast strain** | **Ratio (Low Cu/CM -leu)** | **Ratio (Low Cu -leu/CM -leu)** | **Ratio (Low Cu -leu/Low Cu)** |
|---|---|---|---|
| **W303 (*UPF1*)** | 12.17 (lane 1/lane 3) | 12.53 (± 0.57) (lane 5/lane 3) | 1.03 (± 0.05) (lane 5/lane1) |
| **AAY320 (*upf1*∆)** | 11.48 (lane 2/lane 4) | 12.01 (± 2.78) (lane 6/lane 4) | 1.05 (± 0.24) (lane 6/lane 2) |

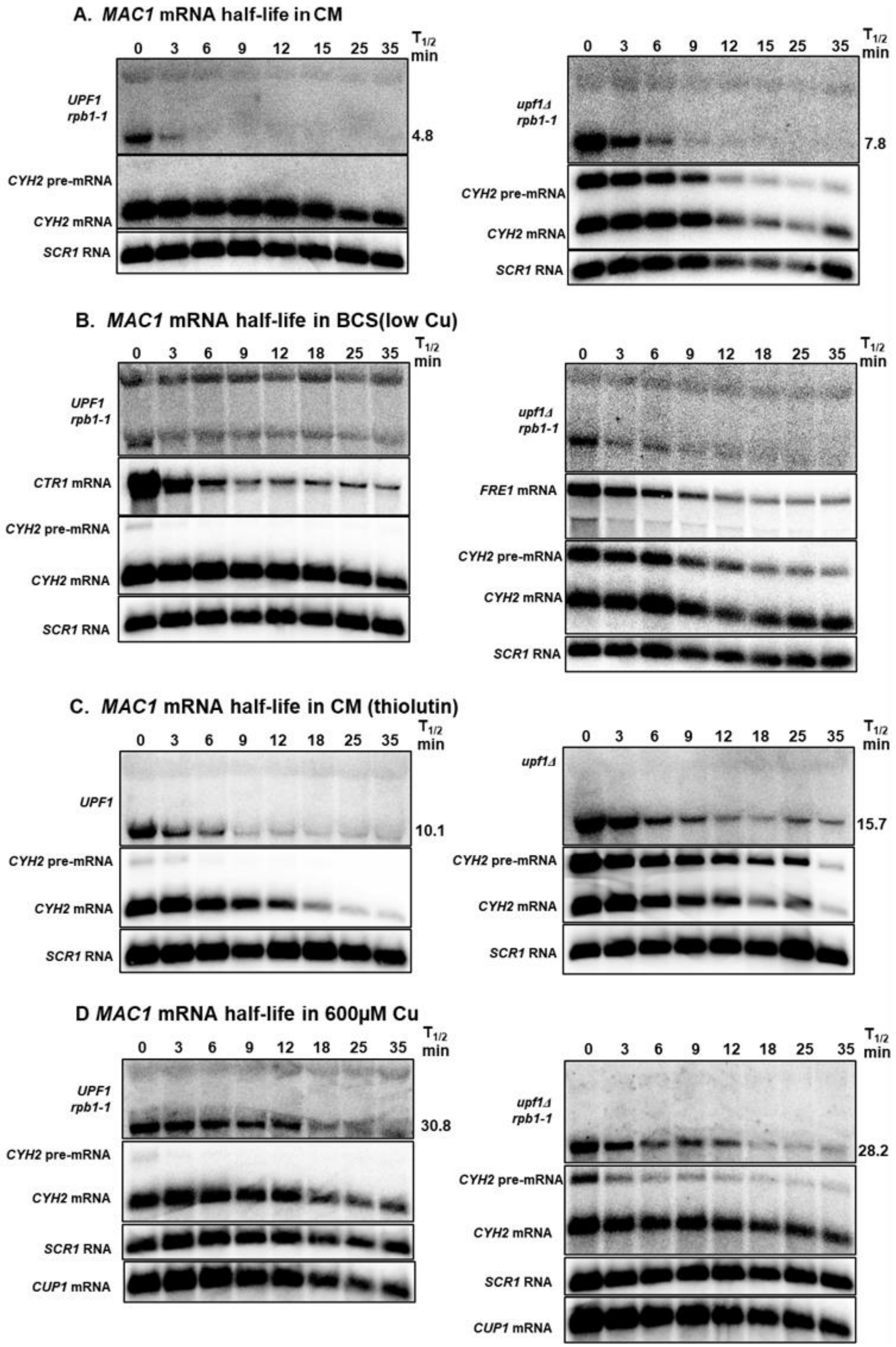

A. MAC1 mRNA half-life in CM
0 3 6 9 12 15 25 35
T1/2 min
UPF1 rpb1-1
4.8
CYH2 pre-mRNA
CYH2 mRNA
SCR1 RNA
upf1Δ rpb1-1
7.8
B. MAC1 mRNA half-life in BCS(low Cu)
0 3 6 9 12 18 25 35
CTR1 mRNA
FRE1 mRNA
C. MAC1 mRNA half-life in CM (thiolutin)
UPF1
10.1
upf1Δ
15.7
D MAC1 mRNA half-life in 600μM Cu
30.8
28.2
CUP1 mRNA

**Supplementary Figure 1.** Representative half-life northern blots of *MAC1* mRNA with controls. The mRNA half-lives were measured with total RNA extracted from wild-type strain AAY334 (*UPF1 rpb1-1*)(1) and NMD mutant strain AAY335 (*upf1Δ rpb1-1*) (1) grown under complete minimal (A), low copper (B) and in 600 µM Cu, high copper (D). mRNA half-lives were measured with total RNA extracted from wild-type strain W303 (*UPF1*) (2), and NMD mutants (*upf1Δ*) (1) grown under complete minimal treated with thiolutin (C). Yeast cells were harvested over a 35-min period at eight time points indicated above the northern blots. The half-lives were determined using SigmaPlot and are shown to the right of the northern blots. All half-life measurements are an average of three independent experiments. *CTR1*, *FRE1*, *CUP1*, *CYH2* and *SCR1* were used as controls. *CTR1* and *FRE1* were used as low copper controls. *CTR1* encodes a high affinity copper transporter of the plasma membrane and is activated under low copper conditions. *FRE1* encodes ferric and cupric reductase, and its expression is induced by low copper and iron levels. *CUP1* was used as a control for high copper because *CUP1* encodes a metallothionein that binds copper. The *CUP1* gene is induced by the Ace1 transcription factor when cells are exposed to elevated copper levels. *CYH2* pre-mRNA was used as an NMD control because *CYH2* pre-mRNA is degraded by NMD. *SCR1* was used as a loading control for all northern blots. *SCR1* is an RNA polymerase III transcript that is not regulated by NMD.

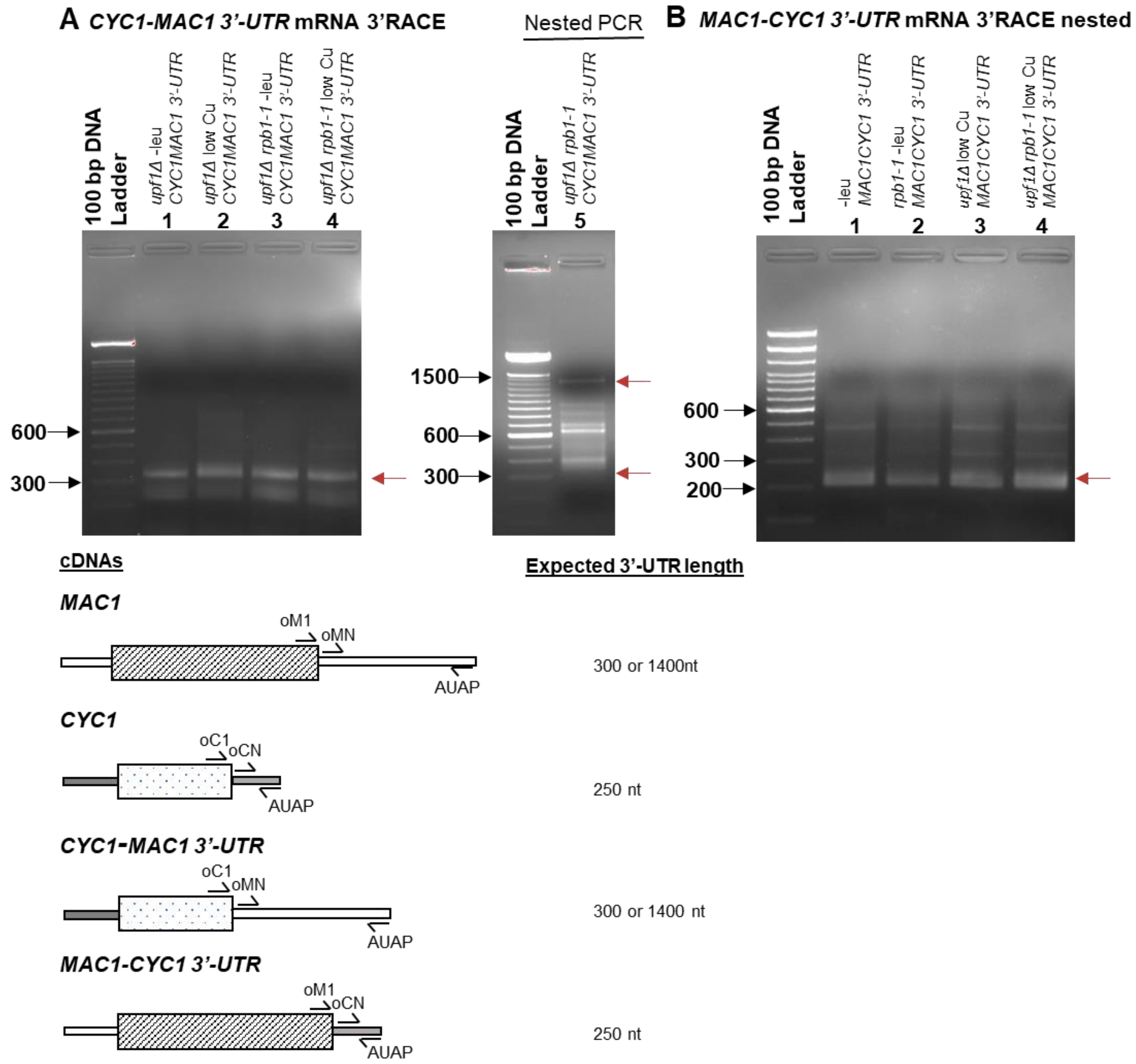


**Supplementary Figure 2.** Precise replacement of the 3′-UTRs in the *CYC1MAC1 3′-UTR* (**A**) and *MAC1CYC1 3′-UTR* (**B**) mRNAs were confirmed by 3′RACE. Schematics of the mRNAs and the primers used in the PCR reactions are shown below the gels. Panel A lanes **1-4** show primary 3′RACE PCR products, while lane 5 show nested PCR products. Lane **1-4**, *CYC1MAC1 3′-UTR* from CM -leu 3′-RACE PCR product using a gene specific primer for the 3′-end of the *CYC1* ORF (oC1) and the Abridged Universal Amplification Primer (AUAP) provided with the 3′RACE kit (Invitrogen Corp. Carlsbad,

CA). The expected band of ~ 300 nt present from 3′-end processing of *CYC1MAC1 3′-UTR* is indicated by an arrow. The additional band shorter band of ~250 nt present corresponding to the endogenous 3′-UTR of *CYC1* mRNA were observed. Additional bands were background because they are present in *MAC1* 3′-UTR nested PCR reactions (Fig. 2D, Lane **2**, *MAC1* mRNA 3′RACE PCR nested with oMN and AUAP primers). Lane **5**, *CYC1-MAC1 3′-UTR* nested PCR products of a reaction using the primary PCR products shown in Lane **3** as template with a primer to the 5′ end of the *MAC1* 3′-UTR (oMN) and the AUAP primer. The expected bands of ~ 300 nt and 1400 nt are indicated by arrows. This data confirms that the *CYC1* 3′-UTR sequences were precisely replaced with the *MAC1* 3′-UTR sequences in the *CYC1MAC1 3′-UTR* construct.

Panel **B**, Lane **1-4**, *MAC1CYC1 3′-UTR* generated using RNA extracted form yeast cells. The primary 3′RACE PCR products were generated with a gene specific primer for the 3′-end of the *MAC1* ORF (oM1) and the Abridged Universal Amplification Primer (AUAP) provided with the 3′RACE kit (Invitrogen Corp. Carlsbad, CA). The primary 3′RACE PCR products were nested with oCN and AUAP primers shows the expected band of ~250 nt (indicated by the arrow). Lane 2, *MAC1CYC1 3′-UTR* generate using RNA extracted form yeast cells grown under from *upf1Δrpb1-1* in low copper -leu nested with oCN and AUAP primers shows the expected band of ~250 nt (indicated by the arrow). This data shows that the *MAC1* 3′-UTR sequences have been precisely replaced by the *CYC1* 3′-UTR sequences in *MAC1CYC1 3′-UTR* construct.

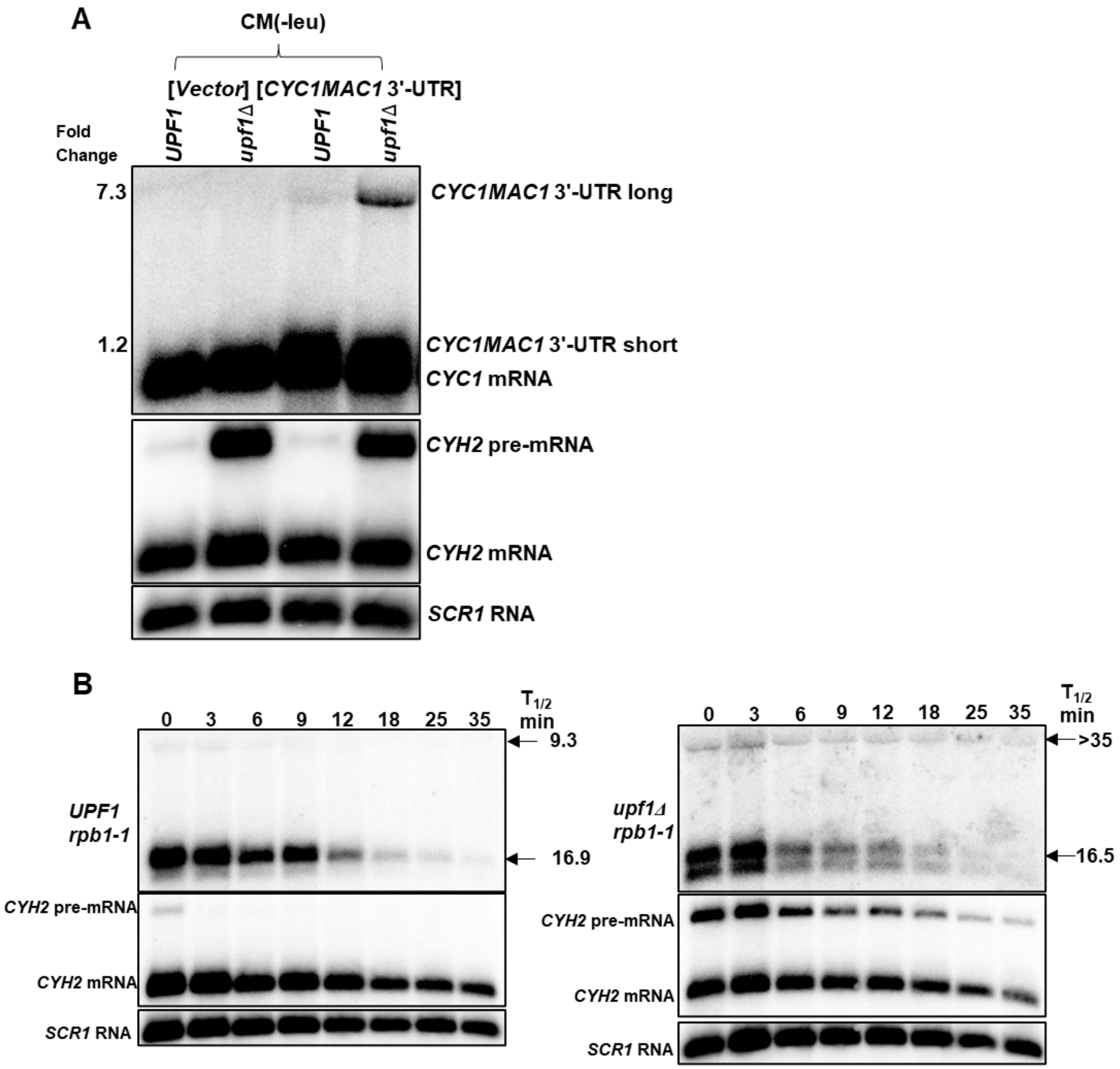


**Supplementary Figure 3.** Steady-state accumulation levels of *CYC1MAC1 3′-UTR* in CM -leu (A). Yeast strains were grown in complete minimal media (CM -leu). The first two lanes of the steady-state northern blot (A) are loaded with RNA from yeast strains lacking the *CYC1MAC1 3′-UTR* mRNA and are transformed with pRS315 (Vector control). The blots were exposed for longer than Fig. 3B to more clearly demonstrate the long *CYC1MAC1* 3′-UTR which is of low abundance. Representative half-life northern blots of *CYC1MAC1 3′-UTR* mRNA with controls in CM -leu (B). All half-life measurements are an average of three independent experiments. *CYH2* and *SCR1* were used as controls as described in Supplementary Figure S1.

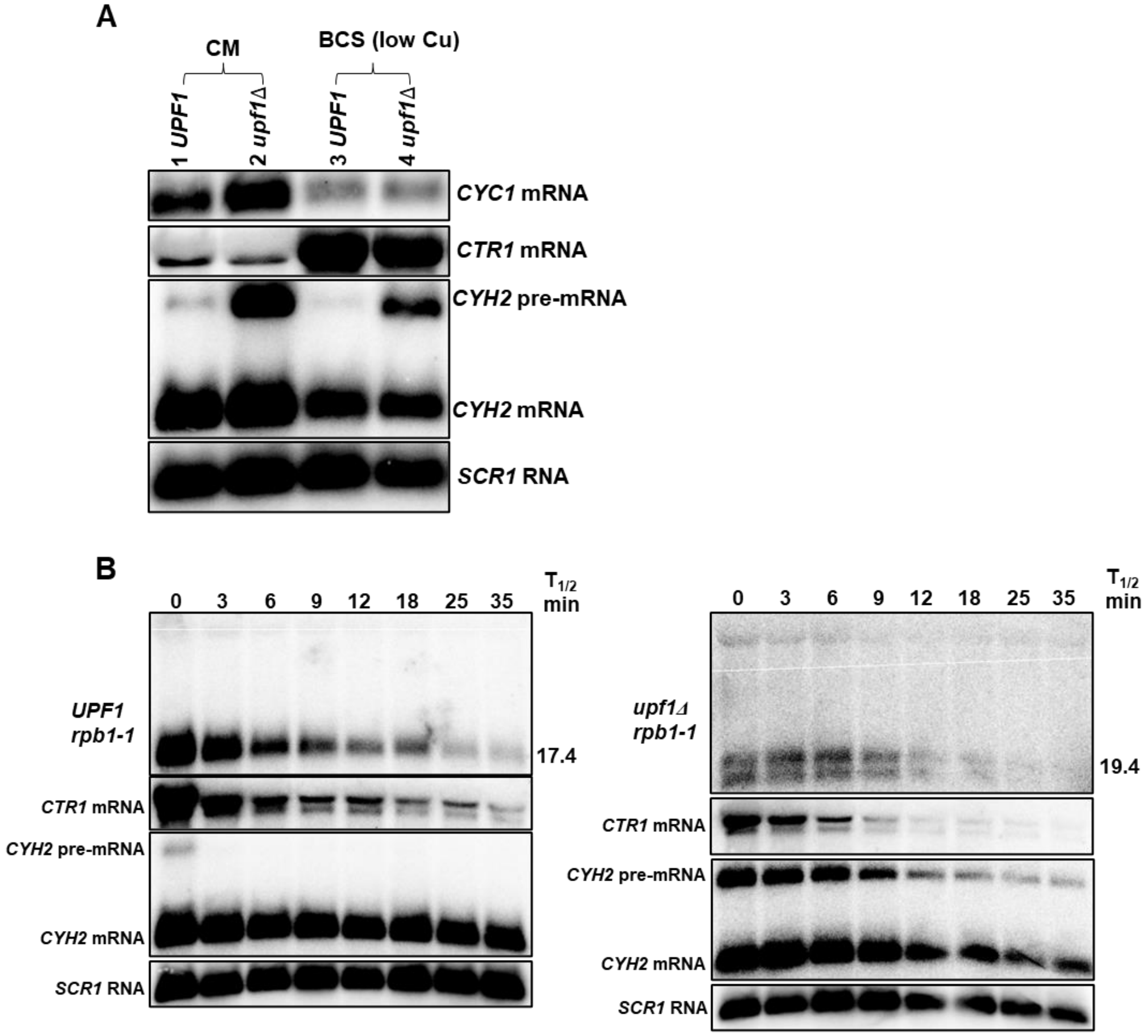


**Supplementary Figure 4.** Steady-state accumulation levels of *CYC1* in CM and low copper (A). Yeast strains were grown in complete minimal (CM) and complete minimal containing 100 µM BCS (low Cu) media. Representative half-life northern blots of *CYC1MAC1 3′-UTR* mRNA with controls in low copper -leu (B). All half-life measurements are an average of three independent experiments. *CTR1*, *CYH2* and *SCR1* were used as controls as described in Supplementary Figure 1.

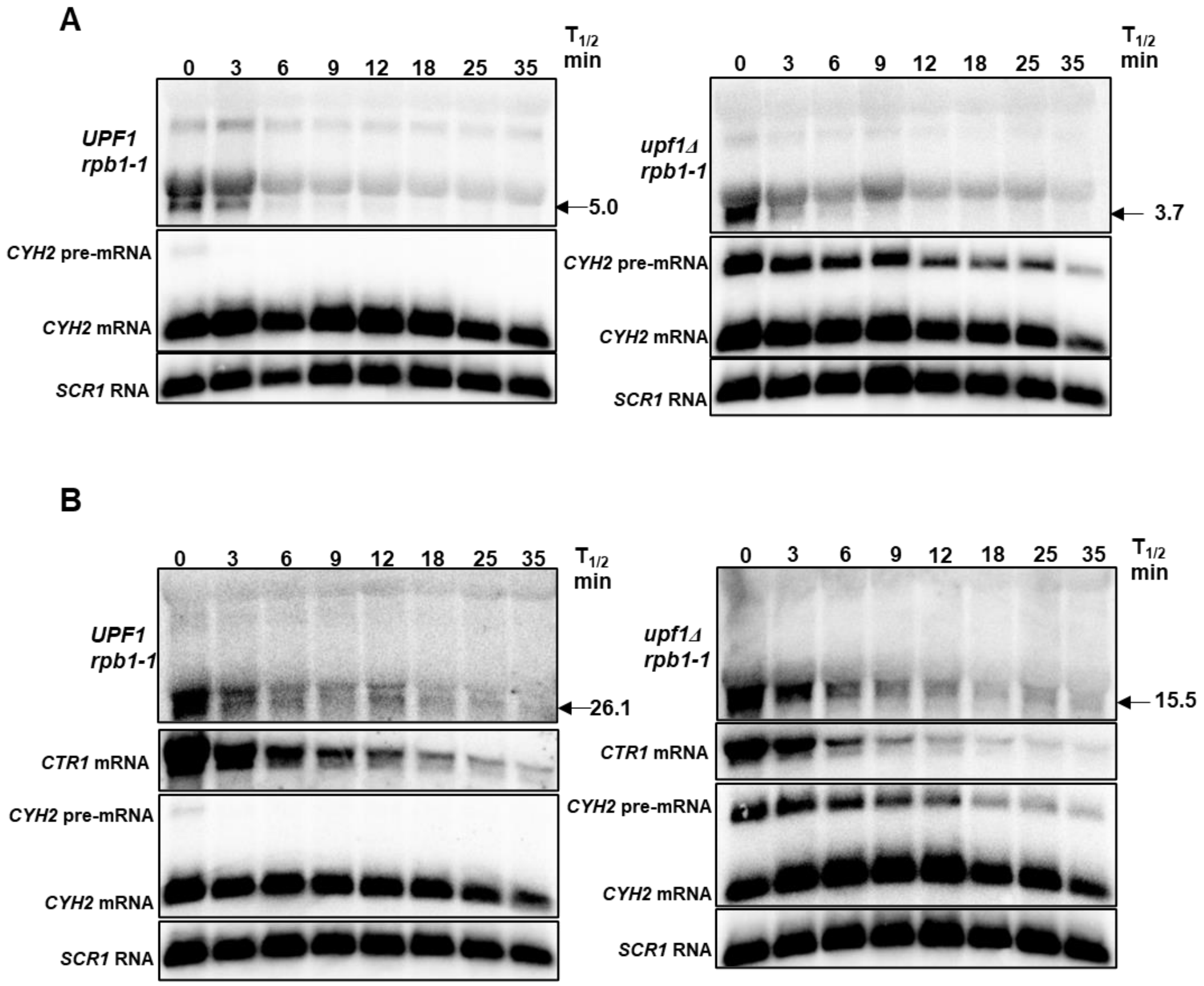


**Supplementary Figure 5.** Representative half-life northern blots of *MAC1CYC1 3'-UTR* mRNA with controls in CM -leu (A) and low copper -leu (B). All half-life measurements are an average of three independent experiments. *CTR1*, *CYH2* and *SCR1* were used as controls as described in Supplementary Figure 1.

Reference

1. Kebaara, B., Nazarenus, T., Taylor, R. and Atkin, A.L. (2003) Genetic background affects relative nonsense mRNA accumulation in wild-type and upf mutant yeast strains. *Curr Genet*, **43**, 171-177.
2. Wente, S.R., Rout, M.P. and Blobel, G. (1992) A new family of yeast nuclear pore complex proteins. *J Cell Biol*, **119**, 705-723.
